# Rationally designed inhibitors of the Musashi protein-RNA interaction by hotspot mimicry

**DOI:** 10.1101/2023.01.09.523326

**Authors:** Nan Bai, Yusuf Adeshina, Igor Bychkov, Yan Xia, Ragul Gowthaman, Sven A. Miller, Abhishek K. Gupta, David K. Johnson, Lan Lan, Erica A. Golemis, Petr B. Makhov, Liang Xu, Manoj M. Pillai, Yanis Boumber, John Karanicolas

## Abstract

RNA-binding proteins (RBPs) are key post-transcriptional regulators of gene expression, and thus underlie many important biological processes. Here, we developed a strategy that entails extracting a “hotspot pharmacophore” from the structure of a protein-RNA complex, to create a template for designing small-molecule inhibitors and for exploring the selectivity of the resulting inhibitors. We demonstrate this approach by designing inhibitors of Musashi proteins MSI1 and MSI2, key regulators of mRNA stability and translation that are upregulated in many cancers. We report this novel series of MSI1/MSI2 inhibitors is specific and active in biochemical, biophysical, and cellular assays. This study extends the paradigm of “hotspots” from protein-protein complexes to protein-RNA complexes, supports the “druggability” of RNA-binding protein surfaces, and represents one of the first rationally-designed inhibitors of non-enzymatic RNA-binding proteins. Owing to its simplicity and generality, we anticipate that this approach may also be used to develop inhibitors of many other RNA-binding proteins; we also consider the prospects of identifying potential off-target interactions by searching for other RBPs that recognize their cognate RNAs using similar interaction geometries. Beyond inhibitors, we also expect that compounds designed using this approach can serve as warheads for new PROTACs that selectively degrade RNA-binding proteins.

## Introduction

RNA-binding proteins (RBPs) play crucial roles in many diverse cellular processes, regulating the life cycle of mRNAs by controlling splicing, polyadenylation, stability, localization and translation, and also modulating the function of non-coding RNAs [1]. Mammalian proteomes are thought to include upwards of 800 RBPs [2,3], corresponding to both RNA-processing enzymes and non-enzymatic RBPs. Given the broad range of functions carried out by RBPs, we sought to devise a general and robust strategy for designing chemical tools that will allow precise manipulation of the interactions between RBPs and their cognate RNAs. Such tools would help unravel the mechanisms of important biological processes controlled by RBPs, and nominate RBPs as feasible targets for therapeutic intervention [4–10].

Despite intense interest, there exist few classes of compounds that target protein-RNA interactions. Efforts to identify such compounds have focused principally on designing assays amenable to high throughput screening [11–13], which indeed has delivered compounds addressing such cancer-relevant RBPs as Musashi-1/2 [14–19], HuR [20–22], eIF4A [23], CELF1 [24], and various splicing factors [25,26]. Among *rationally designed* small-molecule inhibitors that target RBPs, however, all examples reported to date can be categorized into two general classes. The first class comprises nucleoside analogues [27–33], such as anti-HIV-1 NRTIs (nucleoside reverse transcriptase inhibitors), that mimic the chemical structures of natural-occurring nucleosides and rely on enzymatic processing by their targets to form covalent adducts. While nucleoside analogues can be straightforward to design, the inability of these molecules to provide sufficient binding affinity or selectivity without covalent linkage has prevented this strategy from being extended to non-enzymatic RBPs. The second class of compounds comprises allosteric inhibitors [32,34,35], such as anti-HIV-1 NNRTIs (non-nucleoside reverse transcriptase inhibitors), that bind to secondary sites on the protein target and shift its conformation to an inactive state. In principle, allosteric inhibitors could be used to target both enzymatic and non-enzymatic RBPs; in practice, however, challenges associated with both identifying allosteric sites and then finding small molecules to complement these sites has limited the general utility of this approach to all but a few cases. Collectively, the fact that these RNA-binding interfaces are not thought to have evolved to bind small molecules makes them a “non-traditional” class of drug target. Moreover, the relatively flat and polar nature of protein surfaces in this class typically leads to poor performance by structure-based virtual screening (docking) approaches [36]; and given the lack of a known small-molecule binding partners for RBPs, it is even unclear *a priori* that such protein surfaces are suitable for inhibition by any smallmolecule ligand at all [37].

Here, we present a new approach for rationally designing small-molecule inhibitors of RBPs. We draw inspiration from a related class of “non-traditional” drug targets, i.e., protein-protein interfaces. In a protein-protein complex, each of the individual interfacial residues typically do not contribute equally to the energetics of binding; rather, the majority of the binding affinity derives from a small number of “hotspot” residues [38–40]. This observation, in turn, motivated several groups to mimic these key interactions when designing small-molecule inhibitors [41–44]. In this study, we take the “hotspot” paradigm and extend it to protein-RNA interactions.

Our approach entails identifying the chemical moieties of a given RNA that contribute critical interactions to a particular protein-RNA complex, and then identifying small molecules that recapitulate the precise geometrical arrangement of these moieties. Our underlying hypothesis is that compounds capable of mimicking the three-dimensional structure of the RNA “hotspot” will also mimic the energetically dominant interactions in the protein-RNA complex, using a much smaller chemical scaffold. By establishing a new method for reusing these protein-RNA interactions, we circumvent the challenging problem of needing to design interactions that target a flat, polar protein surface.

The approach described above can, in principle, be applied to the structure of any protein-RNA complex. As a first test, we opted to address a target from the most common and well-studied of RNA-binding modules, the RNA-recognition motif (RRM) domain. Hundreds of structures of RRMs have been deposited in the Protein Data Bank, including more than fifty in complex with RNA [45]. Collectively these structures show that RRMs adopt a conserved fold that packs two α-helices against one face of a four-stranded β-sheet; in most cases the opposite face of this β-sheet is then used to bind a single-stranded segment of RNA. Recognition of cognate RNA is usually driven by a cluster of three outward-facing aromatic amino acids on this β-sheet, which often form stacking interactions with a pair of adjacent RNA bases [46]. Accordingly, mutations to the protein that remove these aromatic sidechains have been shown to disrupt binding in representative RRMs [46,47], as has introduction of non-canonical bases to the RNA that alter the pattern of hydrogen bonding groups [48–50]. Despite these shared features, however, the precise geometry of the dinucleotide pair in its complex with the RRM can differ very drastically across members of this family [46].

The Drosophila gene *Musashi (msi*) was originally discovered through its role in sensory organ development [51]; understanding of its role was later expanded to include maintenance of stem cell identity [52]. Mammals encode two close paralogs of *MSI*, known as Musashi-1 (MSI1) and Musashi-2 (MSI2), which also play a role in maintaining stemness [53]. The primary structural elements of Musashi proteins comprise a pair of tandem RRM domains (RRM1 and RRM2) [54], which they use to bind the 3’ UTR region of specific target mRNAs and thus regulate both mRNA stability and translation of the proteins these mRNAs encode. Many of these proteins regulated by MSI1/MSI2 are involved in key signaling pathways that play essential roles in development and are abnormally activated in oncogenesis, such as maintenance of stem cell capacity and promotion of epithelial-mesenchymal transition (NUMB/NOTCH, PTEN/mTOR, TGFβ/SMAD3, MYC) [55,56]; it is thus unsurprising that elevated expression of MSI1/MSI2 has been observed in a broad spectrum of cancers [57–64]. In addition to their roles driving cancer initiation and progression, MSI1/MSI2 have also been implicated in conferring resistance to radiotherapy [65–69], chemotherapy [68,70–72], and targeted therapies [73,74]. Collectively, these observations have positioned MSI1 and MSI2 as tantalizing targets for therapeutic intervention in cancer [55,75].

## Computational Approach

The new computational methods described below are implemented in the Rosetta software suite [76], except where otherwise indicated. Rosetta is freely available for academic use (www.rosettacommons.org), with the new features described here included in the 3.6 release and beyond. Computational methods are summarized below, and then presented in further detail in the *Supporting Methods* section.

### Building “hotspot pharmacophores”

While interfaces between RBPs and their cognate RNAs are mostly flat, complexes involving segments of single-stranded RNA often include a few interfacial nucleobases that are buried much more deeply than the others (**Figure 1a**); this uneven distribution is reminiscent of “hotspot” sidechains in protein-protein complexes [38,39]. The protein has evolved to interact with these buried nucleobases through precise intermolecular aromatic stacking interactions and hydrogen bonding.

**Figure 1:**
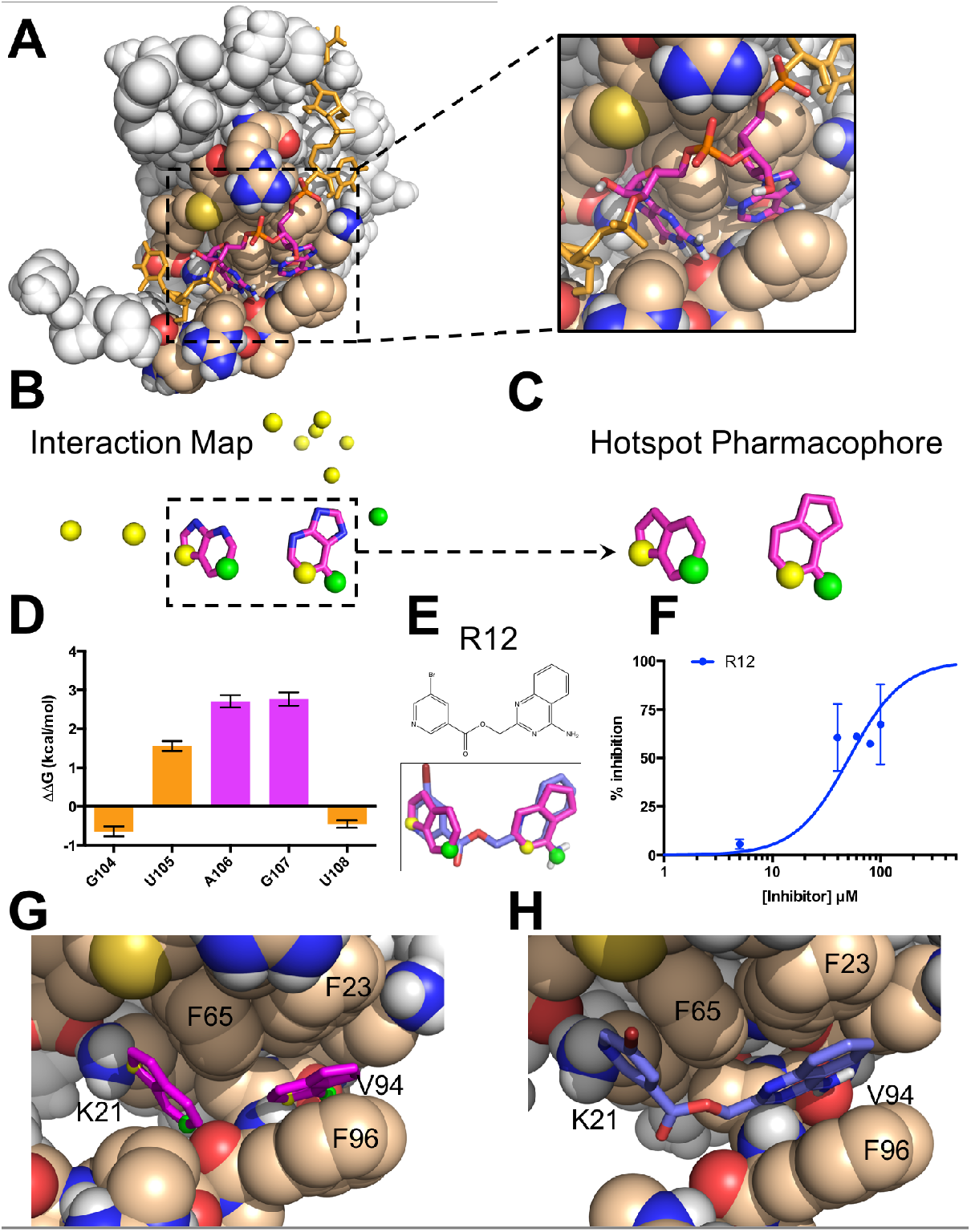
The hotspot mimicry approach. We demonstrate this approach by applying it to the RRM1 domain of Musashi-1. **(A)** The structure of the MSI1 / RNA complex. The RNA (*sticks*) wraps around the protein (*spheres*). Two adjacent bases, A106 and G107 (*magenta*), are buried in a shallow pocket on the protein surface. **(B)** An interaction map is generated from the RNA in the complex, by collecting deeply buried bases (*magenta*) and atoms involved in intermolecular hydrogen bonds (*acceptors shown in yellow, donors in green*). **(C)** Components of the interaction map are clustered in space, and atoms that do not participate in hydrogen bonding are reverted to carbon atoms; this produces a “hotspot pharmacophore.” **(D)** The difference in binding free energy between an RNA harboring a single abasic site versus the original cognate RNA sequence, as determined through competition with a fluorescently-labeled RNA. Positive values indicate diminished binding when a given base is replaced with an abasic site, showing that A106 and G107 contribute more than the other nearby bases to the binding affinity of this interaction. **(E)** The hotspot pharmacophore serves as a template for ligand-based screening, searching for compounds that would mimic the three-dimensional features of the pharmacophore. The screen led to the identification of compound R12, which mimics the geometry of the rings and provides three of the four desired hydrogen bonding groups. **(F)** R12 competes with fluorescein-labeled RNA for MSI1 binding, as observed through a fluorescence polarization assay. These data do not allow the binding affinity to be confidently determined. **(G)** Superposition of the hotspot pharmacophore back onto the protein structure illustrates the interactions that should be captured by an ideal ligand: stacking against three aromatic sidechains, and four intermolecular hydrogen bonds. **(H)** Superposition of R12 onto the protein structure shows that this compound is expected to preserve the aromatic stacking, and to recapitulate three of the four hydrogen bonds.

We have developed an automated framework that distills the structure of a protein-RNA complex to a “hotspot pharmacophore,” which in turn can serve as a template for ligand-based screening. Our framework first picks out those RNA aromatic moieties that are deeply buried in the protein-RNA complex, as well as any RNA atoms involved in intermolecular hydrogen bonds to the protein or ordered water molecules (**Figure 1b**). Any polar atoms on the nucleobases that do not participate in hydrogen bonds are then replaced with carbon atoms, since those polar groups need not be carried forward into inhibitor design. This gives a broad spatial map of the protein-RNA interaction, which typically cannot be spanned by a single drug-like small molecule; we therefore clustered neighboring moieties and advanced each cluster separately. Through this approach, we reduce the structure of the protein-RNA complex to a minimal “hotspot pharmacophore” that encapsulates the key interactions to be recapitulated by a small molecule (**Figure 1c**).

### Identifying complementary ligands

To identify compounds that would mimic RNA’s interactions with the RBP, we then used the hotspot pharmacophore as a template for carrying out ligand-based virtual screening. In order to facilitate rapid characterization of compounds emerging from our screen, we restricted our search to the ~7 million compounds in the ZINC database [77] that are both commercially available, and predicted to have druglike physicochemical properties. We used OMEGA (OpenEye Scientific Software, Santa Fe, NM) [78–80] to build low-energy conformations of each compound, then ROCS (OpenEye Scientific Software, Santa Fe, NM) [81,82] to align each conformation to our hotspot pharmacophore. For each of the top-scoring hits emerging from ROCS, we then used the aligned orientation to position the compound relative to the protein, and evaluated the interaction energy of the protein-ligand complex using the fullatom Rosetta energy function [76].

## Results

### Computational screening against MSI1 RRM1

We applied our “hotspot mimicry” approach to the structure of Musashi-1 RRM1 bound to cognate RNA [54], and found a single hotspot pharmacophore derived from an adjacent pair of buried bases, Adenine106 and Guanine107 (**Figure 1a**). Because no ordered water molecules were included in this NMR structure, the resulting pharmacophore did not include any explicit contribution from solvent. This pharmacophore captures both the aromatic stacking and the hydrogen bonding of the RNA hotspot through its inclusion of ring moieties and donor/acceptor positions, respectively (**Figure 1c**). To test whether these two particular bases indeed serve as a hotspot of the MSI1 RRM1 / RNA interaction, we used a fluorescence polarization (FP) competition assay (see *Supporting Methods*) to measure the binding affinity of RNA variants that maintained the backbone but lacked individual bases. Using this assay, we found that introduction of an abasic site at either of these two positions led to a marked decrease in binding to MSI1 RRM1 (**Figure 1d**). In contrast, introduction of an abasic site at other nearby positions affected binding much less. Confirmation that A106 and G107 serve as hotspot bases of this interaction thus provided experimental evidence supporting the pharmacophore selection from our computational approach.

We then used this pharmacophore as a template for virtual screening. The 12 top-scoring hits identified from the resulting screen could each be classified into one of three diverse chemotypes (**Figure S1**). While none of these scaffolds bear any obvious resemblance in chemical structure to a nucleobase pair, the overlap in three-dimensional shape and hydrogen bonding potential between the hotspot pharmacophore and the modeled conformation of each compound is immediately evident. Despite this strong similarity, none of the 12 hit compounds recapitulated all four of the polar groups included in the hotspot pharmacophore, and only three hit compounds matched to three of the polar groups: R12 (**Figure 1e**), its close analog R4, and R7 (**Figure S2**). Among these three, only R12 showed inhibition in FP competition assay (**Figure 1f**); that said, the binding affinity could not be reasonably quantified because of the low solubility of R12.

As expected, superposition of R21 back onto the hotspot pharmacophore in the context of the protein-RNA complex confirmed that this compound comprised functional groups that would allow recapitulation of the favorable interactions used by the RNA dinucleotide pair. Specifically, the ring moieties in the pharmacophore represent the stacking of nucleobases against Phe23, Phe65, and Phe96 of MSI1, while the hydrogen bonding atoms indicate polar contacts with the sidechain of Lys21 and the backbones of Val94 and Phe96 of MSI1 (**Figure 1g)**. Mimicry of these interactions through the hotspot pharmacophore allows the hit compounds to recapitulate these interactions, as exemplified by R12 (**Figure 1h**). In this model R12 adopts a similar three-dimensional geometry as the hotspot pharmacophore, and thus recapitulates its aromatic stacking and polar interactions (**Figure 1h**).

### Rapid optimization of R12 activity

Guided by our structural model (**Figure 2a**), we then set out to improve potency of R12’s interaction with MSI1. While our initial screen had been restricted to ~7 million compounds in the ZINC database [83], the subsequently-available Enamine database [84] made available (as of 2020) ~8 *billion* compounds: these had not been previously synthesized by Enamine, but were reported to be readily available on-demand. Thus, the Enamine database afforded us an exciting opportunity to carry out “SAR-by-catalog” at a much larger scale than would otherwise have been possible.

**Figure 2:**
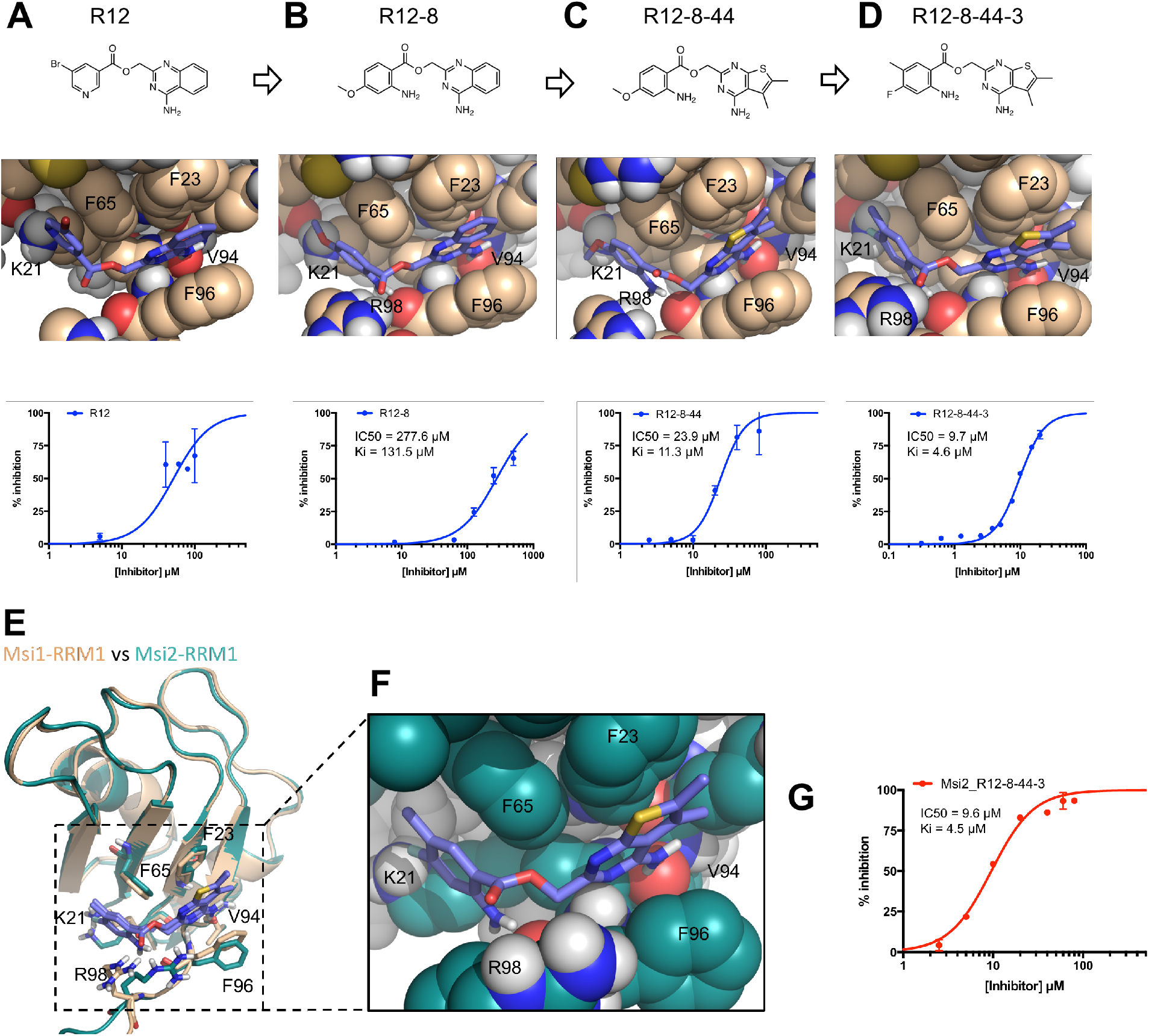
Optimization of R12 to the dual MSI1/MSI2 inhibitor R12-8-44-3. **(A)** R12 was the starting point for optimization, as identified from the computational screen of a limited library; availability of a much larger library was used to enable optimization. **(B)** The left-hand side of R12 was replaced to improve solubility and potency, yielding R12-8. **(C)** The right-hand side of R12-8 was replaced to improve potency, yielding R12-8-44. **(D)** Further exploration of the left-hand side provided improved potency, in R12-8-44-3. **(E)** Superposition of R12-8-44-3 models bound to MSI1 (*wheat*) and MSI2 (*green*). **(F)** Zoomed-in view of the MSI2 model confirms that the expected interactions are unchanged relative to the MSI1 model. **(G)** R12-8-44-3 competes with fluorescein-labeled RNA for

The low solubility of R12 prompted us to begin by looking for alternatives to the bromopyridine group (shown on the left-hand side). Amongst a set of 16 custom analogs that we purchased, R12-7 and R12-8 (**Table S1**), had superior inhibition and solubility than R12. As a secondary validation assay we used Differential Scanning Fluorimetry (DSF) to confirm these compounds’ interaction with the MSI1 RRM1: we found that R12-7 induced noisy changes in the protein’s melting temperature (**Figure S3a**), but that R12-8 led to monotonically increasing stabilization with increasing dose (**Figure S3b**). Out of concern that R12-7’s activity may be associated with compound aggregation, we elected to continue with optimization of R12-8 (**Figure 2b**).

Alignment of R12-8 to our model of MSI1 binding suggested a potential reason for the improved potency. In our initial model of R12, the carbonyl oxygen in its ester linker was positioned in close proximity to the Phe96 backbone carbonyl of MSI1 (**Figure 2a**); beyond simply the lost opportunity for an intermolecular hydrogen bond with the backbone, we expected that there could be electrostatic repulsion between these two negatively charged moieties. By contrast, this ester linker has shifted slightly in our model of R12-8, turning upward to face solvent, and instead engaging MSI1’s backbone carbonyl using R12-8’s newly-added amine (**Figure 2b**). Whereas R12 matched only three of the four desired hotspot pharmacophore features, our model of R12-8 now matched all four (**Figure S3c**).

We next sought to optimize the other side of this compound (the right-hand side), by purchasing another 50 custom analogs from Enamine. While the most potent compound from this second round was R12-8-46, we avoided this compound because catechols are well-known candidate PAINS (pan-assay interference) compounds and can also be oxidized to quinones [85]. Instead, we focused our attention on R12-8-44, which provided more than ten-fold improvement in potency using the FP assay (**Figure 2c**) and a monotonically-increasing melting temperature with increasing ligand concentration (**Figure S4**). R12-8-44 is expected to retain the same polar interactions as R12-8 and R12, but uses slightly different aromatic stacking by replacing R12-8’s 4-quinazolinamine group with 5,6-dimethylthieno[2,3-d]pyrimidin-4-amine.

To further improve potency, we then returned to the left-hand side and assessed seven additional substitutions on the R12-8-44 scaffold (**Figure S5**). Amongst these seven compounds, we found that R12-8-44-3 yielded another 2-fold improvement in potency, ultimately providing an IC_50_ value of 9.7 μM (**Figure 2d**).

### Inhibition of Musashi-2

While expression of MSI1 is tissue-restricted, its homolog MSI2 is ubiquitously expressed [86,87]. Moreover, functional redundancy between the two Musashi family members has led to the proposal that it would be most desirable to have a dual inhibitor that acts on both proteins [88]. Like MSI1, MSI2 includes two RRM domains; the first of these shares 80% sequence identity with MSI1 RRM1. Sequence alignment of MSI1 and MSI2 reveals that all but one of the residues that differ correspond to surface exposed positions far from the hotspot pharmacophore (all but L50M) (**Figure S6**); based on this model, we anticipated that the R12-8-44-3 would also show activity against MSI2.

With this hypothesis in mind, we first built a model of R12-8-44-3 bound to MSI2 by starting from our MSI1-bound model and replacing the 17 residues that differ between MSI1 and MSI2 (**Figure 2e**). The resulting MSI2-bound model (**Figure 2f**) is essentially identical to our earlier MSI1-bound model (**Figure 2d**), implying that R12-8-44-3 should also inhibit MSI2. We tested this using the same FP competition assay and confirmed that R12-8-44-3 indeed inhibits the binding of the MSI2 RRM1 domain to its RNA target with comparable activity as it inhibits MSI1-RRM1 (**Figure 2g**). Thus, we have confirmed that R12-8-44-3 is a dual inhibitor of both MSI1 and MSI2, with the similar potency for each isoform.

**Figure 3:**
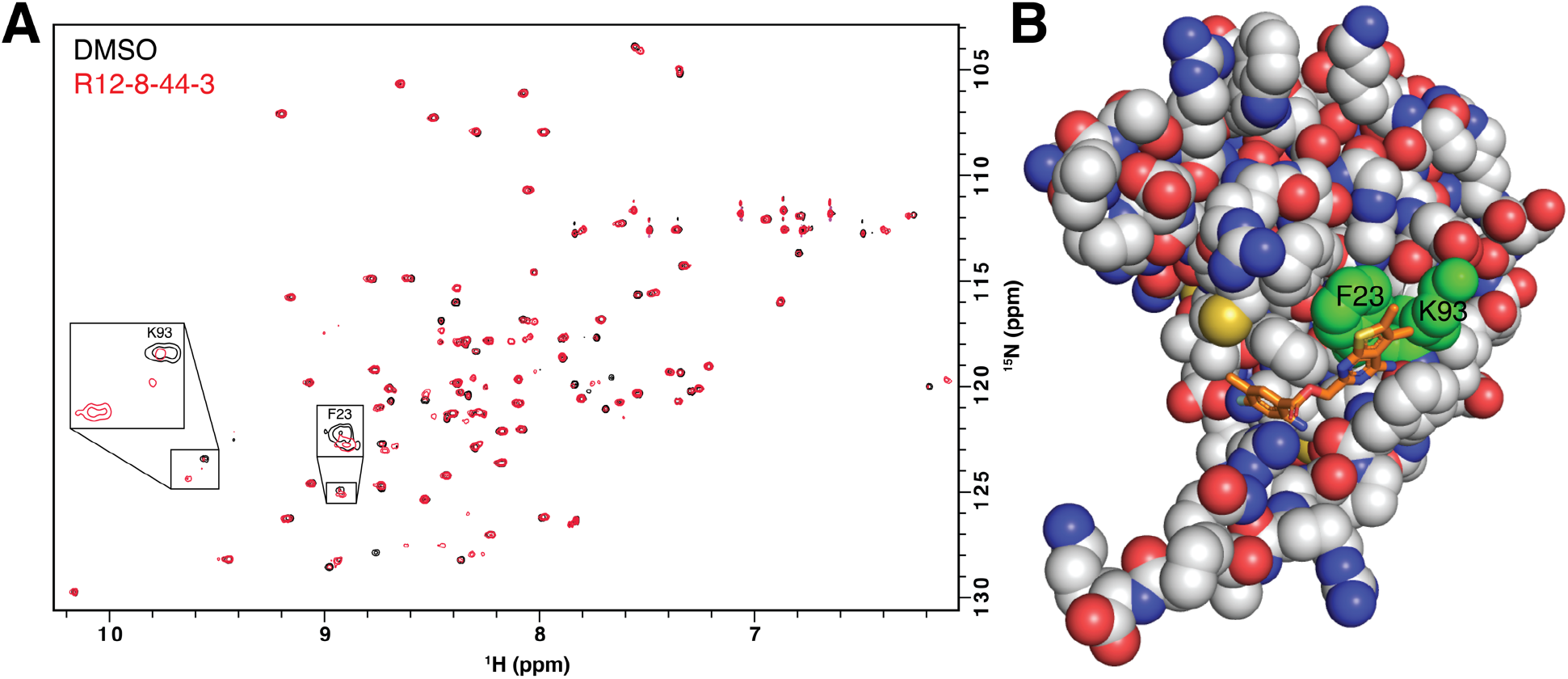
Structural characterization of R12-8-44-3 / MSI1. **(A)** HSQC spectrum of MSI1 RRM1 collected in the presence (*red*) or absence (*black*) of R12-8-44-3. Assigned peaks showing the strongest chemical shift difference are labeled (Phe23, Lys93). **(B)** Mapping the location of these residues to our model of R12-8-44-3/MSI1 confirms that these residues comprise the intended binding site.

### Biophysical characterization of R12-8-44-3

To further characterize R12-8-44-3, we applied differential scanning fluorimetry (DSF) as an orthogonal secondary assay. Surprisingly, addition of R12-8-44-3 did not increase MSI1’s melting temperature as we expected – and as we observed for its parent compounds R12-8 (**Figure S3b**) and R12-8-44 (**Figure S4a**). In fact, it acted in the opposite direction: addition of R12-8-44-3 monotonically decreased MSI1’s melting temperature (**Figure S7**). While this could be a sign of apparent inhibition occurring through aggregation, there are also legitimate inhibitors that reduce their target protein’s melting temperature [89,90]. An alternate explanation could simply be that MSI1’s folding landscape is not simply two-state, and that a partially-folded intermediate stabilized by R12-8-44-3 affects the unfolding transition.

To test the hypothesis that R12-8-44-3 engages MSI1 RRM1 though specific binding interactions (rather than non-specific aggregation), we used HSQC chemical shift mapping. We collected spectra for this construct in the presence and absence of R12-8-44-3 (**Figure 3a**), using previously-reported assignments from a very similar construct [54]. Importantly, we find that only a small number of peaks shift in response to addition of R12-8-44-3: this confirms specific binding to MSI1, rather than non-specific interactions that would imply aggregation. While not all peaks were assigned in our reference spectrum, we gratifyingly find that of the assigned peaks, those with the largest chemical shift differences (Phe23, Lys93) comprise exactly the expected binding site for R12-8-44-3 (**Figure 3b**). Overall, these results strongly support interaction of R12-8-44-3 with the intended binding surface from our computational design.

### Exploring selectivity of R12-8-44-3

Many RRM proteins recognize their target RNAs with high sequence specificity, through additional interactions outside the central RNA dinucleotide [46]. Our mimicry of the MSI1 hotspot was predicated on recapitulating the interactions solely within this dinucleotide; we therefore sought to explore the target selectivity expected for these inhibitors by searching for potential off-target interactions. Starting from every example of protein-RNA complexes in the Protein Data Bank (PDB), we used our computational approach to extract the set of all available hotspot pharmacophores (see *Supporting Methods*). For a given compound of interest, we can then screen all conformers of this molecule against this “library” of 543 unique hotspot pharmacophores (**Figure 4a**). The top-scoring hits in this experiment represent proteins that recognize their cognate RNAs through interaction patterns that can be recapitulated by the compound of interest, making these candidate proteins for off-target binding. In addition to MSI1, there are two other RRM-domain proteins in the PDB that recognize an A-G as the dinucleotide pair: human heterogeneous nuclear ribonucleoprotein A1 (hnRNP A1) [91], and yeast Prp24 [92].

**Figure 4:**
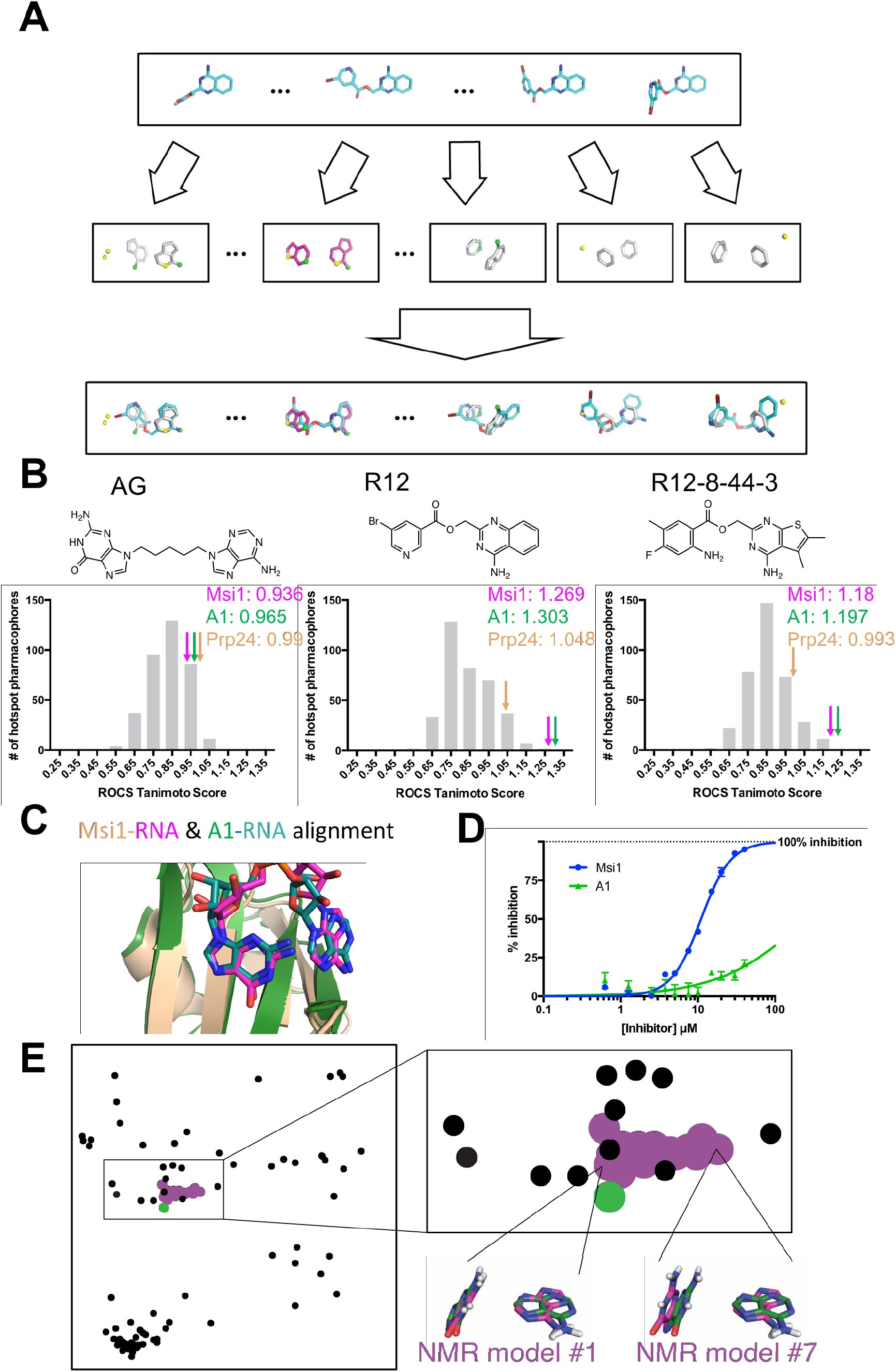
Predicting candidate off-target interactions of a given inhibitor. **(A)** We screened each conformer of a given ligand against the complete set of unique hotspot pharmacophores from other protein-RNA complexes in the PDB. Hits in this screen correspond to other proteins that recognize their cognate RNAs using interaction geometry that can be mimicked by the compound of interest. **(B)** Application of this approach to a hypothetical compound built by connecting adenine and guanine with a flexible linker, to R12, and to R12-8-44-3. High scores correspond to other proteins that recognize their cognate RNAs using interaction geometry that can be mimicked by the compound of interest. The distribution of scores for the complete pharmacophore library is shown, with the score of the MSI1 (*pink arrow*) and hnRNP A1 (*green arrow*) pharmacophores indicated. The artificial compound matches the pharmacophores from many proteins equally well, whereas R12 and R12-8-44-3 match the MSI1 and hnRNP A1 pharmacophores much better than anything else in the library. **(C)** Comparison of the structures of the complexes reveal the basis for identification of hnRNP A1 as a candidate off-target interaction: this protein recognizes its cognate RNA (*green*) with similar positioning of functional groups as MSI1 (*magenta/wheat*), even though the adenine on the right is flipped. **(D)** Evaluation of R12-8-44-3 in an FP competition assay shows that this compound does not inhibit hnRNP A1. **(E)** A projection of the hotspot pharmacophores from all RRM/RNA complexes currently available in the PDB. Each point corresponds to an individual hotspot pharmacophore; this map was generated by using multidimensional scaling analysis to generate the 2D projection that best preserves relative distances between points. With the exception of MSI1, only a single conformation was used for complexes that were solved by NMR. Hotspot pharmacophores from individual members of the MSI1 NMR ensemble are indicated (*magenta*), along with the one from hnRNP A1 (*green*). The compounds described here were identified by computational screening using the hotspot pharmacophore from model #1 of the MSI1 NMR ensemble; this hotspot pharmacophore is very similar to extracted from the hnRNP A1 complex, and accordingly the A-G dinucleotide pair is nearly superposable. In contrast, hotspot pharmacophore from other members of the MSI1 NMR ensemble (such as model #7) are more distant from that of hnRNP A1, and indeed the A-G dinucleotide pair is recognized in a different conformation in these models. The use of highly-distinct models as templates for computational screening may lead to identification of compounds incapable of mimicking the pattern of interactions recognized by other RRM domains, and thus very likely to be selective inhibitors.

We applied this approach first to a hypothetical compound, comprised of adenine and guanine attached by a flexible linker (**Figure 4b**). We built low-energy conformers of this compound, and then evaluated how closely this compound could mimic the three-dimensional geometry of each hotspot pharmacophore found in our library. While this artificial compound can indeed adopt a conformation that aligns well to the MSI1 hotspot pharmacophore (a score of 0.936) and hnRNP A1 (a score of 0.965), this hypothetical A-G compound can also be matched to many other hotspot pharmacophores from the PDB at least as well as these (**Figure 4b**). Thus, this implies that such a compound would bind to many other off-target RBPs, in addition to MSI1 and hnRNP A1. In a sense, this observation underscores the *lack* of selectivity that one might expect from simply mimicking the nucleosides’ chemical structure, without consideration of three-dimensional geometry.

We next carried out the same analysis for our starting compound R12 and optimized compound R12-8-44-3 (**Figure 4b**). By this quantitative analysis (ROCS Tanimoto score), we found that both of these compounds match to the MSI1 hotspot pharmacophore *much* better than they match to any other hotspot pharmacophore extracted from the PDB. This result is unsurprising, given that R12 and R12-8-44-3 lack certain polar groups from the A-G pair (those that did not participate in the MSI1 pharmacophore) and that these compounds have a restricted geometry that allows them only to mimic the specific orientation of the bases needed to complement MSI1 and hnRNP A1.

The reason for R12 and R12-8-44-3 matching well to the hotspot pharmacophore from hnRNP A1 is because of its strong similarity to MSI1’s hotspot pharmacophore: although one of the bases is flipped, the structure of the central RNA dinucleotide in these two complexes is virtually superposable (**Figure 4c**). Overall, this analysis suggests that R12-8-44-3 is likely to be selective for MSI1 over the majority of other RBPs, but that hnRNP A1 could be a potential off-target interaction.

To test this, we expressed and purified hnRNP A1’s RRM1 domain. We confirmed that hnRNP A1 would indeed bind the same fluorescently-labeled RNA used in our previous experiments, and then probed the effect of adding R12-8-44-3 (**Figure 4d**). In this competition experiment, we find that R12-8-44-3 does *not* inhibit hnRNP A1’s RNA binding. We propose that matching to a given hotspot pharmacophore may provide some modest degree of potential binding energy, but that further fine details of the interactions must also be complementary in order to achieve potent binding. Thus, optimization of R12 to R12-8-44-3 enhanced potency for MSI1 by design, and potency for MSI2 because the two proteins are so similar, but would not have impacted the very weak starting affinity for hnRNP A1.

Taking this analysis one step further, we extracted hotspot pharmacophores from each of the 95 RRM/RNA complexes currently present in the PDB and evaluated their similarity in an all-versus-all manner. From these pairwise distances, we then used multidimensional scaling (MDS) analysis to construct the two-dimensional projection that best reflects the pairwise distance between every pair of hotspot pharmacophores: this projection represents a visual “map” of all the hotspot pharmacophores in the PDB (**Figure 4e**).

Unsurprisingly, all of the hotspot pharmacophores built from members of the MSI1 NMR ensemble cluster into a punctate group, reflecting their shared geometric features. The hotspot pharmacophore extracted from the hnRNP A1 crystal structure also overlaps the MSI1 cluster, as expected from the similar recognition of RNA by these two proteins. Though beyond the scope of this study, we expect that screening with a MSI1 template that is more dissimilar to the hnRNP A1 hotspot pharmacophore (such as NMR model #7 instead of model #1), may lead to compounds with even more assurance of selectivity from the outset.

### Cellular activity of R12-8-44-3

A useful secondary outcome of the SAR optimization leading to R12-8-44-3 was the identification of inactive but closely-matched analogs, to serve as negative controls for further study. We therefore selected one such inactive compound and dubbed it “R12-neg” for inclusion in cellular studies. R12-neg retains the substituted thieno[2,3-d]pyrimidine and ester linker from our top active compound R12-8-44-3 (**Figure 5a**), but uses a distinct substitution pattern at the other ring. As a positive control for cellular studies we included the structurally-unrelated neural growth factor (NGF) inhibitor Ro 08-2750 (referred to as “Ro”) [93,94], which has recently been found to inhibit the RNA-binding activity of MSI1/MSI2 *in vitro*, and reduce disease burden using *in vivo* leukemia/lymphoma models [17,95].

**Figure 5:**
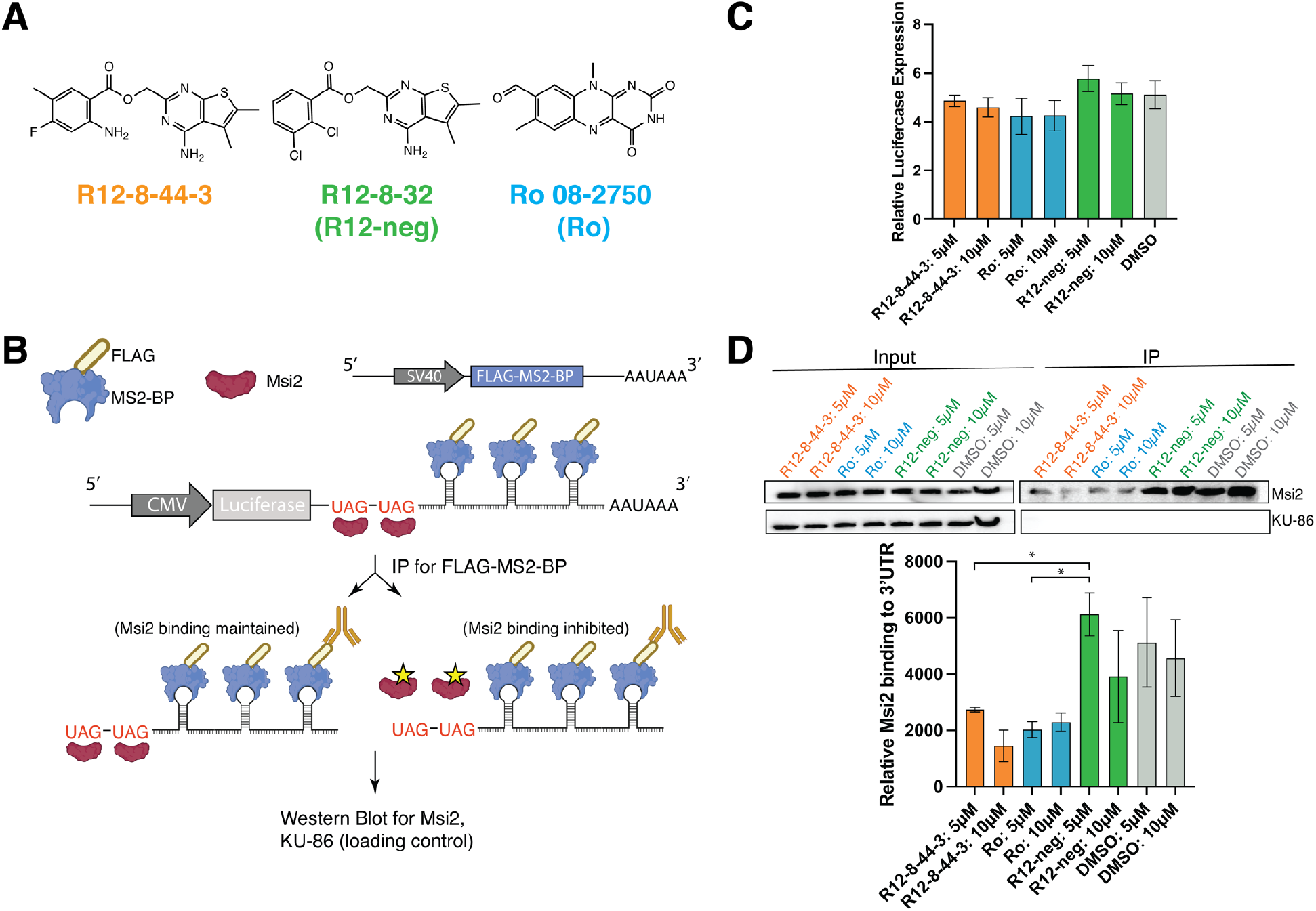
Characterization of R12-8-44-3 in a reporter cell line. **(A)** Chemical structures of R12-8-44-3 and inactive negative control compound R12-8-32 (aka “R12-neg”). As a positive control, Ro 08-2750 (aka “Ro”) is also included in these studies. **(B)** Schematic of reporter assay. HEK293 cells are transfected with a plasmid encoding an arbitrary gene (Luc) with UAG repeats (to recruit MSI2) and MS2 repeats (for pulldown) appended to its 3’ UTR. The same vector also includes FLAG-labeled MS2-BP. Upon anti-FLAG immunoprecipitation (IP), the FLAG-labeled MS2-BP allows purification of the Luc mRNA along with any associated factors. **(C)** Prior to IP, cells express Luc mRNA to a similar extent irrespective of whether they were treated with DMSO, R12-neg, R12-8-44-3, or Ro. **(D)** Prior to IP (“input”), MSI2 is detected in all samples to a similar extent (as is KU-86 loading control). After pulldown of Luc mRNA, MSI2 is found associated with the mRNA in cells treated with DMSO or R12-neg, but not in cells that had been treated with R12-8-44-3 or Ro. Representative image is shown; quantification is plotted as the average of 2 independent experiments ± S.E.M. Statistical analysis was performed using unpaired two tailed t-test, with * indicating p<0.05. Uncropped Western blots from this experiment are provided as **Figure S8**.

To determine whether inhibition of MSI1/MSI2 by R12-8-44-3 would apply in a cellular context, we adapted an aptamer pull-down reporter assay that we previously used to validate candidate Musashi target genes [96]. Briefly, our approach is a variation of the MS2-TRAP (MS2-tagged RNA affinity purification) method, which uses bacteriophage protein MS2-BP to extract from cells any factors that associate with RNAs containing a specific stem-loop structure (MS2) from the phage genome [97]. Here, we designed an mRNA construct that would include a series of MS2 hairpins and also a MSI2 recognition sequence. We transfected this construct into cells that also express FLAG-tagged MS2-BP (a protein unrelated to MSI2). By affinity purification using the FLAG tag, proteins bound to the designed mRNA can be extracted from cells.

To test for inhibition of MSI2 in cells, we transfected HEK293 cells (with a high baseline MSI2 expression), with the pMS2-LUC-3’UTR from our previous studies [96]. This vector appends onto the 3’-UTR of an arbitrary gene (in this case luciferase) two key elements: UAG-motifs comprising a MSI2 recognition sequence, and a series of five MS2 hairpins. The same vector also encodes the FLAG-MS2-BP fusion needed for affinity purification (**Figure 5b**). Cells were then treated with each compound of interest for 24 h, and then the cells were subsequently harvested and used for anti-FLAG immunoprecipitation. While luciferase was used as the arbitrary gene for MSI2 recognition in this experiment, we did not explicitly monitor luciferase activity in these studies. Using qPCR we evaluated the amount of Luc mRNA in cells treated with R12-8-44-3, R12-neg, or Ro, and found this to be unchanged (**Figure 5c**).

Upon treatment with R12-8-44-3, R12-neg, or Ro, we also found that the total amount of MSI2 in cells was unaffected (**Figure 5d**). Affinity purification of the reporter mRNA for cells treated with DMSO control confirms MSI2 binding to this mRNA. For cells treated with R12-8-44-3 (or Ro), however, much less MSI2 is purified from cells using this reporter mRNA (**Figure 5d**): beyond its activity in biochemical assays, this result suggests that R12-8-44-3 also inhibits MSI2 in cells and prevents engagement with its target genes. Consistent with results from its earlier biochemical characterization, R12-neg does not disrupt MSI2 pulldown.

To explore the effect of these MSI2 inhibitors in a cancer-relevant cellular context, we selected two MSI2-dependent cancer cell lines: non-small cell lung cancer cell line PC9 and leukemia cell line K562 [73,98–101]. Upon treatment of PC9 cells with Ro, R12-8-44-3, or negative control R12-neg, we found at multiple time points that cellular MSI2 protein levels were unchanged (**Figure 6a**). To confirm target engagement, we therefore adapted the cellular thermal shift assay (CETSA) [102]. To determine experimental conditions, we first harvested cells and incubated at increasing temperatures; we then removed aggregated protein by centrifugation and used Western blotting to probe the amount of MSI2 remaining in the soluble fraction (**Figure 6b, Figure S9**). Having established 49 °C as the midpoint of the transition under these (non-equilibrium) conditions, we then treated cells with our candidate MSI2 inhibitor for 3 h, and heated to 49 °C. In both PC9 cells (**Figure 6c**) and K562 cells (**Figure 6d**), we find that R12-8-44-3 confers protection against thermal-induced aggregation, confirming its direct interaction with MSI2 in both cell lines.

**Figure 6:**
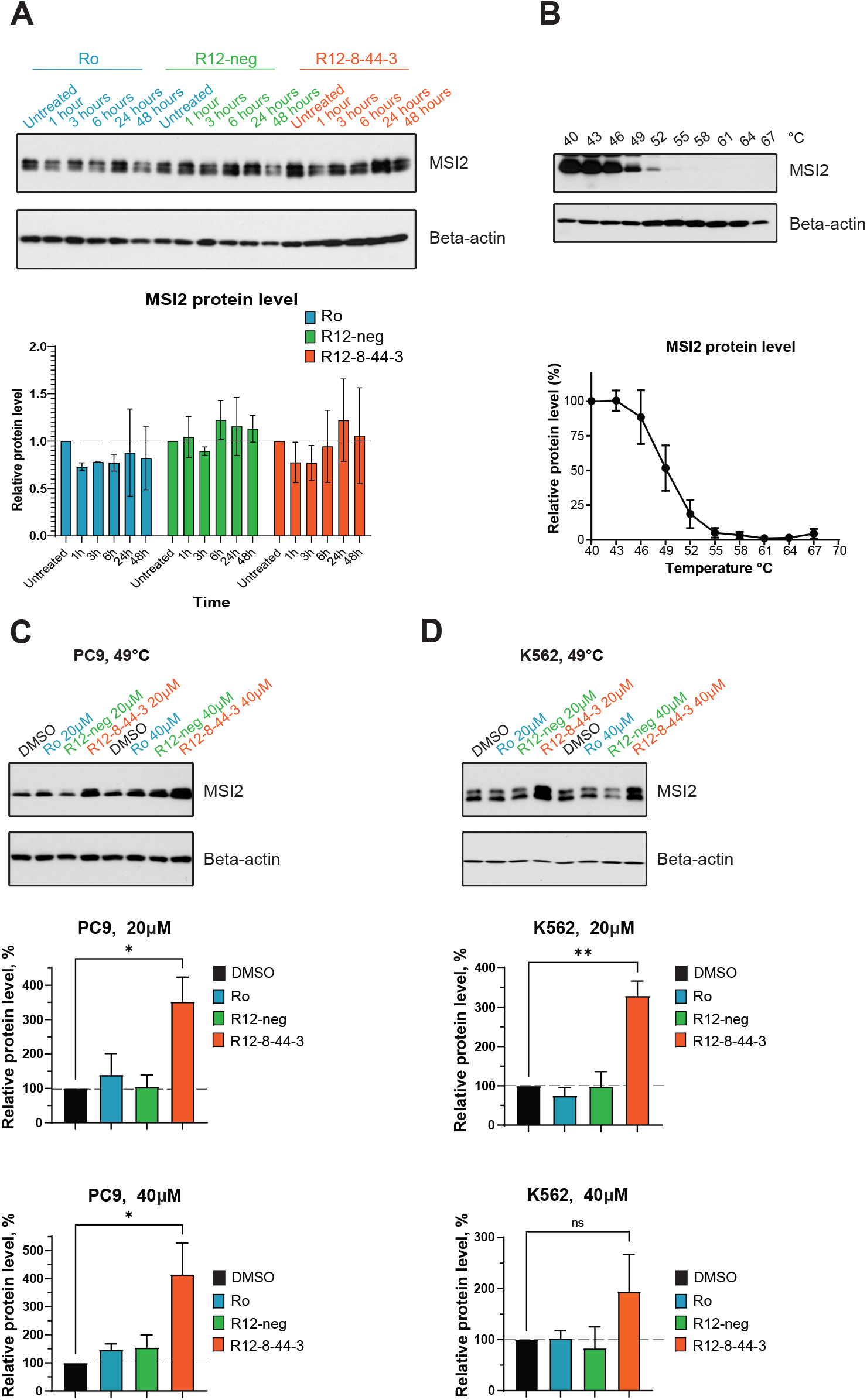
Direct cellular engagement of MSI2 interaction by R12-44-3. **(A)** Immunoblot from PC9 cells treated with Ro (positive control), R12-neg (negative control), or R12-8-44-3. Cells were treated with 20 μM compound for 1, 3, 6, 24, or 48 hours. MSI2 protein abundance was then quantified and normalized to untreated sample. One representative image is shown; quantification is presented as the average ± S.E.M. from 3 independent replicates. **(B)** Thermal-induced aggregation of cellular MSI2 in the PC9 cell line. Cells were briefly incubated at temperatures between 40 °C to 67 °C, then centrifuged and the amount of MSI2 remaining in the soluble fraction was quantified by Western blotting. Data are presented as the average ± S.E.M. from 3 independent replicates, normalized first to beta-actin and then to the data at 40 °C. **(C)** PC9 cells were treated for 3 hours with 20 μM or 40 μM of the indicated compounds, then heated to 49 °C and lysed. One representative image is shown; quantification is presented as the average ± S.E.M. from 3 independent replicates, normalized first to beta-actin and then to DMSO control. **(D)** K562 cells were treated for 3 hours with 20 μM or 40 μM of the indicated compounds, then heated to 49 °C and lysed. One representative image is shown; quantification is presented as the average ± S.E.M. from 3 independent replicates, normalized first to beta-actin and then to DMSO control. Statistical analysis was performed using unpaired two tailed t-test, with * indicating p<0.05, and ** indicating p<0.01.

Having shown that R12-8-44-3 engages MSI2 in these two cell lines, we next explored whether treatment with this compound also inhibits the interaction of MSI2 with known target mRNAs. Among previously described targets of MSI2 are *PTP4A, SMAD3, EGFR*, and *MSI2* itself [73,98–101]. To test for disruption of MSI2 with these cognate mRNAs upon treatment with R12-8-44-3, we used mRNA immunoprecipitation (RIP): we used an anti-MSI2 antibody to immunoprecipitate MSI2, then reverse transcribed and analyzed by qPCR to determine the amount of each specific mRNA that was bound to MSI2. As an important control to ensure that the resulting mRNA was indeed brought down by MSI2, we additionally verified that using a negative control antibody (instead of anti-MSI2) brought with it 100-fold less of each target mRNA (**Figure S10**). As a separate negative control, GAPDH mRNA was also quantified in the context of these studies (since GAPDH is not thought to be a target of MSI2).

PC9 cells were then treated with 10 μM of each compound for 8 hours and then analyzed by RIP. For each MSI2 target gene, treatment with R12-8-44-3 (but not R12-neg) led to a statistically significant reduction in the amount of mRNA associated with MSI2 (**Figure 7a**). Collectively then, these results suggest that R12-8-44-3 inhibits binding of MSI2 to its mRNA targets in cells.

**Figure 7:**
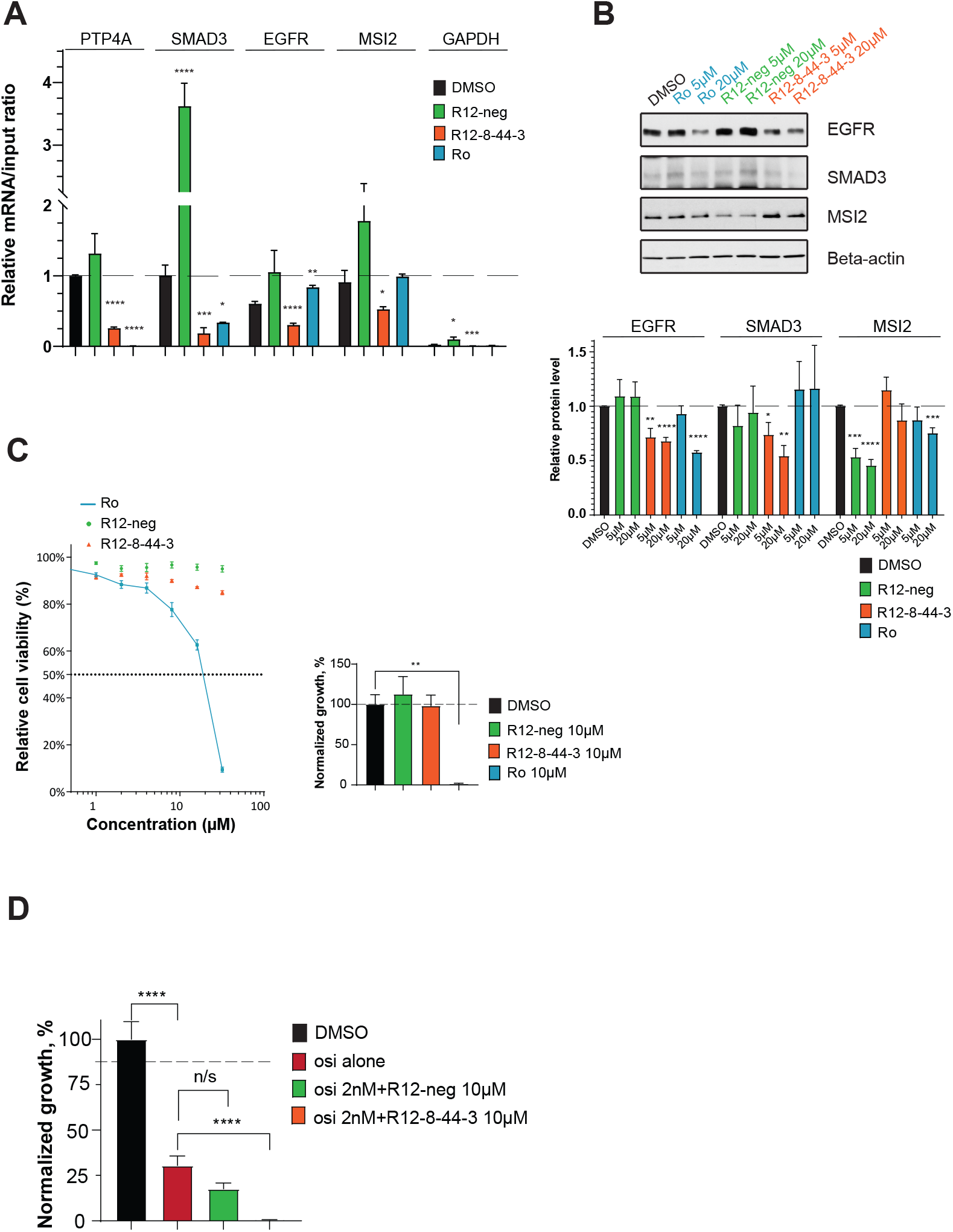
Cellular characterization of MSI2 inhibition by R12-8-44-3. **(A)** Quantification of mRNA immunoprecipitation (RIP) results. Cells were treated with 10 μM of the indicated compounds for 8 hours. After immunoprecipitation with anti-MSI2, mRNA targets were quantified. The amount of each target mRNA in the pulldown was calculated relative to the input, and results from all targets were normalized to *PTP4A1*. Treating cells with R12-8-44-3 disrupted the interaction with all four cognate mRNAs (*PTP4A, SMAD3, EGFR*, and *MSI2*). Data are presented as the average ± S.E.M. from at least 3 independent replicates. **(B)** Immunoblot from PC9 cells treated with Ro (positive control), R12-neg (negative control), or R12-8-44-3. Cells were treated with indicated compounds at two concentrations (5 μM and 20 μM) for 48 hours, then protein levels of relevant MSI2 targets were determined (EGFR, SMAD3, and MSI2). One representative image is shown; quantification is presented as the average ± S.E.M. from at least 3 independent replicates, normalized first to beta-actin and then to DMSO-treated cells. **(C)** Cell viability dose-response quantified by CellTiter-Blue after 3 days (*left*) and cell proliferation with 10μM of compounds quantified by clonogenic assay after 7 days (*right*). Each is normalized to DMSO control, and presented as the average ± S.E.M. from 3 independent replicates. **(D)** Cell viability quantified by clonogenic assay of PC9 cell line after 7-day treatment with 2 nM osimertinib (osi alone), osimertinib with 10 μM R12-8-44-3, or osimertinib with 10 μM R12-neg. Data are normalized to DMSO, and are presented as the average ± S.E.M. from 9 independent replicates. Statistical analysis was performed using unpaired two tailed t-test. In all plots, * indicates p<0.05, ** indicates p<0.01, *** indicates p<0.001, and **** indicates p<0.0001.

Binding of Musashi proteins to specific target mRNAs can lead to either up- or down-regulation of the corresponding protein product [55]. MSI2 depletion (by RNA knockdown) in PC9 cells leads to reduced SMAD3 and EGFR, prompting us to explore whether pharmacological inhibition of MSI2 confers the same downstream effect. Consistent with the intended mechanism of MSI2 inhibition, we found that treating PC9 cells with either 5 μM or 20 μM R12-8-44-3 (but not R12-neg) phenocopied the effect of MSI2 depletion on these two targets, and it did so without changing the amount of MSI2 (**Figure 7b**). Similar effects were also observed in K562 cells (**Figure S11a**).

Effects of MSI2 on downstream targets can be readily probed using RNA knockdown to deplete MSI2; however, the cellular dependence on MSI2 activity for cell growth/survival is more challenging to evaluate. Development of R12-8-44-3 thus affords us with a tool to directly evaluate the consequence of MSI2 pharmacological inhibition in cells. Accordingly, we treated PC9 cells with R12-8-44-3 and used CellTiter-Blue assay to probe cell viability after 3 days and a clonogenic assay to probe cell proliferation after 7 days.

To our surprise, we found that MSI2 inhibition by R12-8-44-3 had no effect on cell growth/survival in either assay (**Figure 7c**). This observation stands in stark contrast to Ro, which dramatically diminishes cell viability in these experiments. We also extended these studies to include K562 cells (**Figure S11b**), and we found results in alignment with our observations from PC9 cells. It is difficult to determine precisely why Ro and R12-8-44-3 have such dramatically different behavior in this assay: one possibility is that Ro, a repurposed compound originally reported for activity unrelated to MSI2, may exert its cytotoxic effect in these cell lines through molecular targets unrelated to MSI2. As raised by others recently [18], our results call into question whether MSI2 inhibition alone may be insufficient as a therapeutic strategy; certainly though, these results suggest that MSI2 may be required for survival only in certain cancer cells and not others.

With respect to non-small cell lung cancer (NSCLC) specifically, overexpression of MSI2 has been identified as a mechanism underlying resistance to inhibitors of EGFR – and more importantly, depletion of MSI2 by RNA knockdown increased sensitivity to the third-generation EGFR inhibitor osimertinib both *in vitro* and *in vivo* [73,74]. Despite lack of sensitivity to R12-8-44-3 alone, we therefore hypothesized that R12-8-44-3 would sensitize PC9 cells (an EGFR-mutant lung cancer cell line) to osimertinib. We first used CellTiter-Blue assay to evaluate the response of PC9 cells to a 3-day treatment of osimertinib, either alone or in combination with 10 μM R12-8-44-3: this experiment showed a small but statistically significant decrease in cell viability due to inclusion of R12-8-44-3 (**Figure S12**). Further, clonogenic assays performed over 7 days demonstrated that pharmacological inhibition of MSI2 provides a robust combined effect with osimertinib (**Figure 7d**).

In summary, we conclude that R-12-8-44-3 directly inhibits binding of MSI2 to target mRNAs in human lung cancer and leukemic cell line models. While this inhibition alone does not prove cytotoxic, at least in these particular cell lines, MSI2 inhibition nonetheless sensitizes EGFR-mutant lung cancer cells to the EGFR inhibitor osimertinib. Taken together, these data demonstrate that R12-8-44-3 directly binds and inhibits MSI2 biologic activity in cellular models, and that it can serve as a starting point for identifying potentially useful synergistic combinations with other approved targeted therapies.

## Discussion

The ability to rationally design selective inhibitors of RNA-binding proteins in a robust and general way will enable development of new tool compounds to help elucidate cellular processes mediated by these interactions. Naturally-occurring examples have shown that proteins can mimic certain structural features of RNAs [103,104]; here, we instead encode a key RNA epitope on a small-molecule scaffold. We demonstrate the application of our approach using Musashi-1 and Musashi-2, leading to a novel class of inhibitors that disrupt the RNA-binding activity of these tumor-promoting proteins. By using the hotspot pharmacophore as a template for ligand-based screening, our approach circumvents the challenge of explicitly designing *de novo* interactions against a relatively flat and polar protein surface.

The major advantages of this mimicry approach are its generality and simplicity. In our first application of this RNA mimicry approach, we elected to restrict our initial screening to commercially available compounds. Though none of the resulting hit compounds provided complete recapitulation of the desired hotspot interactions, we found that one of these, R12, complemented the protein surface without steric clashes and provided a starting point for new inhibitors of MSI1, thus validating the computational approach. In parallel with this work, a contemporaneous study also sought to design inhibitors of RBPs via mimicry of the cognate RNA [105]. This study reports inhibitors of a different RRM-domain protein, HuR, by manually selecting key moieties from the RNA to mimic, then using computational approaches to design compounds accessible through multicomponent reaction chemistry. While STD-NMR confirmed the interaction of some of these compounds with the protein, the binding affinities for these compounds were not reported.

To optimize R12, we then drew from an enormous library of make-on-demand compounds to carry out a modern form of SAR-by-catalog. The availability of this resource allowed us to rapidly drive forward optimization, and ultimately led us to an inhibitor with single-digit micromolar binding affinity, R12-8-44-3. Moving forward, we expect that ongoing increases in the size of this (and competitors’) “virtual catalogs” will greatly facilitate medicinal chemistry optimization for many other projects as well.

With regards to selectivity, further experimental evidence will be necessary to thoroughly and explicitly determine whether the compounds reported here engage in unanticipated interactions with any other RBPs. Nonetheless, the computational approach described herein also provides a potential strategy to identify likely off-target interaction partners: we propose that aligning candidate compounds to hotspot pharmacophores extracted from other RBPs can help identify potential off-target interactions. This can allow prioritization of select off-target RBPs for explicit biochemical testing, rather than simply collecting arbitrary off-target RBPs for evaluation. In this vein, we were especially pleased to note that R12-8-44-3 showed no inhibition for the RBP predicted as its most-likely off-target interaction, hnRNP A1.

We do note, however, that predictions of potential off-target interactions in this manner are necessarily limited: both by the incompleteness of the set of RBP complexes in the PDB, and by the fact that complexes solved using x-ray crystallography are present as single points on this map, instead of clusters that reflect conformational flexibility. Even with this limitation, however, already the utility of this approach to identify potential off-target interactions is clear. While we cannot explicitly confirm that R12-8-44-3 does not inhibit any other RBPs in the human proteome, we can at least provide a rationale for why selectivity should be expected from this compound. Looking ahead, this general strategy may also enable predicting the potential selectivity of a given compound at an earlier stage, which in turn may help prioritize specific scaffolds and drive further focused optimization. Further, the most “isolated” points on our map of pharmacophore space (**Figure 4e**) may represent the most distinctive hotspot pharmacophores in the PDB: the uniqueness of these protein targets may make them particularly amenable to design of highly selective inhibitors. We also note that the utility of our approach is also likely to be dramatically enriched in the near future by emerging structure-prediction methods using deep learning, to allow near-atomic resolution predictions for protein-RNA complexes [106]: these new tools will certainly broaden the scope of RBPs that can be targeted using the strategy presented here, and may also allow richer and more impactful selectivity predictions.

Finally, we do acknowledge that our optimization of the R12 series did not lead to extremely potent compounds; we suspect that this may be an intrinsic limitation of the relatively flat binding site available on the protein surface. That said, PROTACs (PROteolysis TArgeting Chimeras) [107,108] have emerged as a viable strategy for addressing challenging targets, and may be exquisitely well suited for advancing these compounds. In considering development of RNA-mimicking inhibitors as warheads for development of new PROTACs, we note specifically that the binding affinity for the target has proven not to be a major determinant of effective target degradation. Thus, even if achieving highly potent direct inhibitors of RRM domains remains challenging, selective inhibitors may nonetheless offer a path forward for unlocking the tantalizing biology of RBPs, both as novel chemical probes and also as potential starting points for new therapeutics.

## Methods

Detailed descriptions of computational and experimental methods are provided in the *Supporting Methods* section.

## Acknowledgements

This work used the Extreme Science and Engineering Discovery Environment (XSEDE) allocation MCB130049, which is supported by National Science Foundation grant number ACI-1548562. Y.B. was also supported by the 2022 Translational Bridge Award from Northwestern University. This work was supported by grants from NIGMS (R01GM123336 and R01GM141513) and grants from the NCI (R01CA178831, R01CA218802, and R01CA218802). This research was also funded in part through the NIH/NCI Cancer Center Support Grants P30CA006927 and P30CA060553.

## References

1. Muller-McNicoll M, Neugebauer KM. How cells get the message: dynamic assembly and function of mRNA-protein complexes. Nat Rev Genet. 2013; 14:275–87.

2. Castello A, Fischer B, Eichelbaum K, Horos R, Beckmann BM, Strein C, Davey NE, Humphreys DT, Preiss T, Steinmetz LM, Krijgsveld J, Hentze MW. Insights into RNA biology from an atlas of mammalian mRNA-binding proteins. Cell. 2012; 149:1393–406.

3. Baltz AG, Munschauer M, Schwanhausser B, Vasile A, Murakawa Y, Schueler M, Youngs N, Penfold-Brown D, Drew K, Milek M, Wyler E, Bonneau R, Selbach M, Dieterich C, Landthaler M. The mRNA-bound proteome and its global occupancy profile on protein-coding transcripts. Mol Cell. 2012; 46:674–90.

4. Pascale A, Govoni S. The complex world of post-transcriptional mechanisms: is their deregulation a common link for diseases? Focus on ELAV-like RNA-binding proteins. Cell Mol Life Sci. 2012; 69:501–17.

5. Kapeli K, Yeo GW. Genome-wide approaches to dissect the roles of RNA binding proteins in translational control: implications for neurological diseases. Front Neurosci. 2012; 6:144.

6. Khalil AM, Rinn JL. RNA-protein interactions in human health and disease. Semin Cell Dev Biol. 2011; 22:359–65.

7. Mohibi S, Chen X, Zhang J. Cancer the ‘RBP’eutics-RNA-binding proteins as therapeutic targets for cancer. Pharmacol Ther. 2019; 203:107390.

8. Elcheva IA, Spiegelman VS. Targeting RNA-binding proteins in acute and chronic leukemia. Leukemia. 2021; 35:360–76.

9. Holmqvist E, Vogel J. RNA-binding proteins in bacteria. Nat Rev Microbiol. 2018; 16:601–15.

10. Mittal N, Roy N, Babu MM, Janga SC. Dissecting the expression dynamics of RNA-binding proteins in posttranscriptional regulatory networks. Proc Natl Acad Sci US A. 2009; 106:20300–5.

11. Byun WG, Lim D, Park SB. Small-molecule modulators of protein-RNA interactions. Curr Opin Chem Biol. 2022; 68:102149.

12. D’Agostino VG, Sighel D, Zucal C, Bonomo I, Micaelli M, Lolli G, Provenzani A, Quattrone A, Adami V. Screening Approaches for Targeting Ribonucleoprotein Complexes: A New Dimension for Drug Discovery. SLAS Discov. 2019; 24:314–31.

13. Julio AR, Backus KM. New approaches to target RNA binding proteins. Curr Opin Chem Biol. 2021; 62:13–23.

14. Lan L, Appelman C, Smith AR, Yu J, Larsen S, Marquez RT, Liu H, Wu X, Gao P, Roy A, Anbanandam A, Gowthaman R, Karanicolas J, De Guzman RN, Rogers S, Aube J, Ji M, Cohen RS, Neufeld KL, Xu L. Natural product (-)-gossypol inhibits colon cancer cell growth by targeting RNA-binding protein Musashi-1. Mol Oncol. 2015; 9:1406–20.

15. Lan L, Liu H, Smith AR, Appelman C, Yu J, Larsen S, Marquez RT, Wu X, Liu FY, Gao P, Gowthaman R, Karanicolas J, De Guzman RN, Rogers S, Aube J, Neufeld KL, Xu L. Natural product derivative Gossypolone inhibits Musashi family of RNA-binding proteins. BMC Cancer. 2018; 18:809.

16. Lan L, Liu J, Xing M, Smith AR, Wang J, Wu X, Appelman C, Li K, Roy A, Gowthaman R, Karanicolas J, Somoza AD, Wang CCC, Miao Y, De Guzman R, Oakley BR, Neufeld KL, Xu L. Identification and Validation of an Aspergillus nidulans Secondary Metabolite Derivative as an Inhibitor of the Musashi-RNA Interaction. Cancers (Basel). 2020; 12.

17. Minuesa G, Albanese SK, Xie W, Kazansky Y, Worroll D, Chow A, Schurer A, Park SM, Rotsides CZ, Taggart J, Rizzi A, Naden LN, Chou T, Gourkanti S, Cappel D, Passarelli MC, Fairchild L, Adura C, Glickman JF, Schulman J, Famulare C, Patel M, Eibl JK, Ross GM, Bhattacharya S, Tan DS, Leslie CS, Beuming T, Patel DJ, Goldgur Y, Chodera JD, Kharas MG. Small-molecule targeting of MUSASHI RNA-binding activity in acute myeloid leukemia. Nat Commun. 2019; 10:2691.

18. Zhang X, Su K, Liu Y, Zhu D, Pan Y, Ke X, Qu Y. Small Molecule Palmatine Targeting Musashi-2 in Colorectal Cancer. Front Pharmacol. 2021; 12:793449.

19. Clingman CC, Deveau LM, Hay SA, Genga RM, Shandilya SM, Massi F, Ryder SP. Allosteric inhibition of a stem cell RNA-binding protein by an intermediary metabolite. Elife. 2014; 3.

20. Wu X, Xu L. The RNA-binding protein HuR in human cancer: A friend or foe? Adv Drug Deliv Rev. 2022; 184:114179.

21. Majumder M, Chakraborty P, Mohan S, Mehrotra S, Palanisamy V. HuR as a molecular target for cancer therapeutics and immune-related disorders. Adv Drug Deliv Rev. 2022; 188:114442.

22. Della Volpe S, Linciano P, Listro R, Tumminelli E, Amadio M, Bonomo I, Elgaher WAM, Adam S, Hirsch AKH, Boeckler FM, Vasile F, Rossi D, Collina S. Identification of N,N-arylalkyl-picolinamide derivatives targeting the RNA-binding protein HuR, by combining biophysical fragment-screening and molecular hybridization. Bioorg Chem. 2021; 116:105305.

23. Zerio CJ, Cunningham TA, Tulino AS, Alimusa EA, Buckley TM, Moore KT, Dodson M, Wilson NC, Ambrose AJ, Shi T, Sivinski J, Essegian DJ, Zhang DD, Schurer SC, Schatz JH, Chapman E. Discovery of an eIF4A Inhibitor with a Novel Mechanism of Action. J Med Chem. 2021; 64:15727–46.

24. Tan Y, Sun X, Xu Y, Tang B, Xu S, Lu D, Ye Y, Luo X, Diao X, Li F, Wang T, Chen J, Xu Q, Wu X. Small molecule targeting CELF1 RNA-binding activity to control HSC activation and liver fibrosis. Nucleic Acids Res. 2022; 50:2440–51.

25. Tang J, Xie Y, Huang J, Zhang L, Jiang W, Li Z, Bian J. A critical update on the strategies towards small molecule inhibitors targeting Serine/arginine-rich (SR) proteins and Serine/arginine-rich proteins related kinases in alternative splicing. Bioorg Med Chem. 2022; 70:116921.

26. Zhang Y, Wang M, Meng F, Yang M, Chen Y, Guo X, Wang W, Zhu Y, Guo Y, Feng C, Tian S, Zhang H, Li H, Sun J, Wang W. A novel SRSF3 inhibitor, SFI003, exerts anticancer activity against colorectal cancer by modulating the SRSF3/DHCR24/ROS axis. Cell Death Discov. 2022; 8:238.

27. Squires KE. An introduction to nucleoside and nucleotide analogues. Antivir Ther. 2001; 6 Suppl 3:1–14.

28. Cihlar T, Ray AS. Nucleoside and nucleotide HIV reverse transcriptase inhibitors: 25 years after zidovudine. Antiviral Res. 2010; 85:39–58.

29. Bitterman PB, Polunovsky VA. Attacking a nexus of the oncogenic circuitry by reversing aberrant eIF4F-mediated translation. Mol Cancer Ther. 2012; 11:1051–61.

30. Menendez-Arias L, Alvarez M, Pacheco B. Nucleoside/nucleotide analog inhibitors of hepatitis B virus polymerase: mechanism of action and resistance. Curr Opin Virol. 2014; 8C:1–9.

31. James SH, Prichard MN. Current and future therapies for herpes simplex virus infections: mechanism of action and drug resistance. Curr Opin Virol. 2014; 8C:54–61.

32. Das K, Arnold E. HIV-1 reverse transcriptase and antiviral drug resistance. Part 1. Curr Opin Virol. 2013; 3:111–8.

33. Ewald B, Sampath D, Plunkett W. Nucleoside analogs: molecular mechanisms signaling cell death. Oncogene. 2008; 27:6522–37.

34. Biswas S, Sukla S, Field HJ. Helicase-primase inhibitors for herpes simplex virus: looking to the future of non-nucleoside inhibitors for treating herpes virus infections. Future Med Chem. 2014; 6:45–55.

35. Das K, Arnold E. HIV-1 reverse transcriptase and antiviral drug resistance. Part 2. Curr Opin Virol. 2013; 3:119–28.

36. Gowthaman R, Deeds EJ, Karanicolas J. Structural properties of non-traditional drug targets present new challenges for virtual screening. J Chem Inf Model. 2013; 53:2073–81.

37. Fauman EB, Rai BK, Huang ES. Structure-based druggability assessment--identifying suitable targets for small molecule therapeutics. Curr Opin Chem Biol. 2011; 15:463–8.

38. Clackson T, Wells JA. A hot spot of binding energy in a hormone-receptor interface. Science. 1995; 267:383–6.

39. Moreira IS, Fernandes PA, Ramos MJ. Hot spots--a review of the protein-protein interface determinant amino-acid residues. Proteins. 2007; 68:803–12.

40. Rajamani D, Thiel S, Vajda S, Camacho CJ. Anchor residues in protein-protein interactions. Proc Natl Acad Sci U S A. 2004; 101:11287–92.

41. Koes DR, Camacho CJ. Small-molecule inhibitor starting points learned from protein-protein interaction inhibitor structure. Bioinformatics. 2012; 28:784–91.

42. Thanos CD, DeLano WL, Wells JA. Hot-spot mimicry of a cytokine receptor by a small molecule. Proc Natl Acad Sci U S A. 2006; 103:15422–7.

43. Christ F, Voet A, Marchand A, Nicolet S, Desimmie BA, Marchand D, Bardiot D, Van der Veken NJ, Van Remoortel B, Strelkov SV, De Maeyer M, Chaltin P, Debyser Z. Rational design of smallmolecule inhibitors of the LEDGF/p75-integrase interaction and HIV replication. Nat Chem Biol. 2010; 6:442–8.

44. Liu S, Wu S, Jiang S. HIV entry inhibitors targeting gp41: from polypeptides to small-molecule compounds. Curr Pharm Des. 2007; 13:143–62.

45. Daubner GM, Clery A, Allain FH. RRM-RNA recognition: NMR or crystallography...and new findings. Curr Opin Struct Biol. 2013; 23:100–8.

46. Maris C, Dominguez C, Allain FH. The RNA recognition motif, a plastic RNA-binding platform to regulate post-transcriptional gene expression. FEBS J. 2005; 272:2118–31.

47. Auweter SD, Oberstrass FC, Allain FH. Sequence-specific binding of single-stranded RNA: is there a code for recognition? Nucleic Acids Res. 2006; 34:4943–59.

48. Nolan SJS, J. C.; Tuite, J. B.; Cecere, K. L.; Baranger, A. M. Recognition of an essential adenine at a protein-RNA interface: comparison of the contribution of hydrogen bonds and a stacking interaction. J Am Chem Soc. 1999:2.

49. Benitex Y, Baranger AM. Recognition of essential purines by the U1A protein. BMC Biochem. 2007; 8:22.

50. Tuite JB, Shiels JC, Baranger AM. Substitution of an essential adenine in the U1A-RNA complex with a non-polar isostere. Nucleic Acids Res. 2002; 30:5269–75.

51. Nakamura M, Okano H, Blendy JA, Montell C. Musashi, a neural RNA-binding protein required for Drosophila adult external sensory organ development. Neuron. 1994; 13:67–81.

52. Siddall NA, McLaughlin EA, Marriner NL, Hime GR. The RNA-binding protein Musashi is required intrinsically to maintain stem cell identity. Proc Natl Acad Sci U S A. 2006; 103:8402–7.

53. Sutherland JM, McLaughlin EA, Hime GR, Siddall NA. The Musashi family of RNA binding proteins: master regulators of multiple stem cell populations. Adv Exp Med Biol. 2013; 786:233–45.

54. Ohyama T, Nagata T, Tsuda K, Kobayashi N, Imai T, Okano H, Yamazaki T, Katahira M. Structure of Musashi1 in a complex with target RNA: the role of aromatic stacking interactions. Nucleic Acids Res. 2012; 40:3218–31.

55. Kudinov AE, Karanicolas J, Golemis EA, Boumber Y. Musashi RNA-Binding Proteins as Cancer Drivers and Novel Therapeutic Targets. Clin Cancer Res. 2017; 23:2143–53.

56. Bley N, Hmedat A, Muller S, Rolnik R, Rausch A, Lederer M, Huttelmaier S. Musashi-1-A Stemness RBP for Cancer Therapy? Biology (Basel). 2021; 10.

57. Fan LF, Dong WG, Jiang CQ, Xia D, Liao F, Yu QF. Expression of putative stem cell genes Musashi-1 and beta1-integrin in human colorectal adenomas and adenocarcinomas. Int J Colorectal Dis. 2010; 25:17–23.

58. Ma YH, Mentlein R, Knerlich F, Kruse ML, Mehdorn HM, Held-Feindt J. Expression of stem cell markers in human astrocytomas of different WHO grades. J Neurooncol. 2008; 86:31–45.

59. Seigel GM, Hackam AS, Ganguly A, Mandell LM, Gonzalez-Fernandez F. Human embryonic and neuronal stem cell markers in retinoblastoma. Mol Vis. 2007; 13:823–32.

60. Toda M, Iizuka Y, Yu W, Imai T, Ikeda E, Yoshida K, Kawase T, Kawakami Y, Okano H, Uyemura K. Expression of the neural RNA-binding protein Musashi1 in human gliomas. Glia. 2001; 34:1–7.

61. Wang XY, Penalva LO, Yuan H, Linnoila RI, Lu J, Okano H, Glazer RI. Musashi1 regulates breast tumor cell proliferation and is a prognostic indicator of poor survival. Mol Cancer. 2010; 9:221.

62. Ye F, Zhou C, Cheng Q, Shen J, Chen H. Stem-cell-abundant proteins Nanog, Nucleostemin and Musashi1 are highly expressed in malignant cervical epithelial cells. BMC Cancer. 2008; 8:108.

63. Yokota N, Mainprize TG, Taylor MD, Kohata T, Loreto M, Ueda S, Dura W, Grajkowska W, Kuo JS, Rutka JT. Identification of differentially expressed and developmentally regulated genes in medulloblastoma using suppression subtraction hybridization. Oncogene. 2004; 23:3444–53.

64. Kameda-Smith MM, Zhu H, Luo EC, Suk Y, Xella A, Yee B, Chokshi C, Xing S, Tan F, Fox RG, Adile AA, Bakhshinyan D, Brown K, Gwynne WD, Subapanditha M, Miletic P, Picard D, Burns I, Moffat J, Paruch K, Fleming A, Hope K, Provias JP, Remke M, Lu Y, Reya T, Venugopal C, Reimand J, Wechsler-Reya RJ, Yeo GW, Singh SK. Characterization of an RNA binding protein interactome reveals a context-specific post-transcriptional landscape of MYC-amplified medulloblastoma. Nat Commun. 2022; 13:7506.

65. de Araujo PR, Gorthi A, da Silva AE, Tonapi SS, Vo DT, Burns SC, Qiao M, Uren PJ, Yuan ZM, Bishop AJ, Penalva LO. Musashi1 Impacts Radio-Resistance in Glioblastoma by Controlling DNA-Protein Kinase Catalytic Subunit. Am J Pathol. 2016; 186:2271–8.

66. Troschel FM, Palenta H, Borrmann K, Heshe K, Hua SH, Yip GW, Kiesel L, Eich HT, Gotte M, Greve B. Knockdown of the prognostic cancer stem cell marker Musashi-1 decreases radio-resistance while enhancing apoptosis in hormone receptor-positive breast cancer cells via p21(WAF1/CIP1). J Cancer Res Clin Oncol. 2021; 147:3299–312.

67. Falke I, Troschel FM, Palenta H, Loblein MT, Bruggemann K, Borrmann K, Eich HT, Gotte M, Greve B. Knockdown of the stem cell marker Musashi-1 inhibits endometrial cancer growth and sensitizes cells to radiation. Stem Cell Res Ther. 2022; 13:212.

68. Loblein MT, Falke I, Eich HT, Greve B, Gotte M, Troschel FM. Dual Knockdown of Musashi RNA-Binding Proteins MSI-1 and MSI-2 Attenuates Putative Cancer Stem Cell Characteristics and Therapy Resistance in Ovarian Cancer Cells. Int J Mol Sci. 2021; 22.

69. Lin JC, Tsai JT, Chao TY, Ma HI, Chien CS, Liu WH. MSI1 associates glioblastoma radioresistance via homologous recombination repair, tumor invasion and cancer stem-like cell properties. Radiother Oncol. 2018; 129:352–63.

70. Lee J, An S, Choi YM, Lee J, Ahn KJ, Lee JH, Kim TJ, An IS, Bae S. Musashi-2 is a novel regulator of paclitaxel sensitivity in ovarian cancer cells. Int J Oncol. 2016; 49:1945–52.

71. Han Y, Ye A, Zhang Y, Cai Z, Wang W, Sun L, Jiang S, Wu J, Yu K, Zhang S. Musashi-2 Silencing Exerts Potent Activity against Acute Myeloid Leukemia and Enhances Chemosensitivity to Daunorubicin. PLoS One. 2015; 10:e0136484.

72. Liu F, Yang H, Zhang X, Sun X, Zhou J, Li Y, Liu Y, Zhuang Z, Wang G. Inhibition of Musashi-1 enhances chemotherapeutic sensitivity in gastric cancer patient-derived xenografts. Exp Biol Med (Maywood). 2022; 247:868–79.

73. Makhov P, Bychkov I, Faezov B, Deneka A, Kudinov A, Nicolas E, Brebion R, Avril E, Cai KQ, Kharin LV, Voloshin M, Frantsiyants E, Karnaukhov N, Kit OI, Topchu I, Fazliyeva R, Nikonova AS, Serebriiskii IG, Borghaei H, Edelman M, Dulaimi E, Golemis EA, Boumber Y. Musashi-2 (MSI2) regulates epidermal growth factor receptor (EGFR) expression and response to EGFR inhibitors in EGFR-mutated non-small cell lung cancer (NSCLC). Oncogenesis. 2021; 10: 29.

74. Yiming R, Takeuchi Y, Nishimura T, Li M, Wang Y, Meguro-Horike M, Kohno T, Horike SI, Nakata A, Gotoh N. MUSASHI-2 confers resistance to third-generation EGFR-tyrosine kinase inhibitor osimertinib in lung adenocarcinoma. Cancer Sci. 2021; 112:3810–21.

75. Palacios F, Yan XJ, Ferrer G, Chen SS, Vergani S, Yang X, Gardner J, Barrientos JC, Rock P, Burack R, Kolitz JE, Allen SL, Kharas MG, Abdel-Wahab O, Rai KR, Chiorazzi N. Musashi 2 influences chronic lymphocytic leukemia cell survival and growth making it a potential therapeutic target. Leukemia. 2021; 35:1037–52.

76. Leaver-Fay A, Tyka M, Lewis SM, Lange OF, Thompson J, Jacak R, Kaufman K, Renfrew PD, Smith CA, Sheffler W, Davis IW, Cooper S, Treuille A, Mandell DJ, Richter F, Ban YE, Fleishman SJ, Corn JE, Kim DE, Lyskov S, Berrondo M, Mentzer S, Popovic Z, Havranek JJ, Karanicolas J, Das R, Meiler J, Kortemme T, Gray JJ, Kuhlman B, Baker D, Bradley P. ROSETTA3: an object-oriented software suite for the simulation and design of macromolecules. Methods Enzymol. 2011; 487:545–74.

77. Irwin JJ, Sterling T, Mysinger MM, Bolstad ES, Coleman RG. ZINC: A Free Tool to Discover Chemistry for Biology. J Chem Inf Model. 2012.

78. Hawkins PC, Skillman AG, Warren GL, Ellingson BA, Stahl MT. Conformer generation with OMEGA: algorithm and validation using high quality structures from the Protein Databank and Cambridge Structural Database. J Chem Inf Model. 2010; 50:572–84.

79. Hawkins PC, Nicholls A. Conformer generation with OMEGA: learning from the data set and the analysis of failures. J Chem Inf Model. 2012; 52:2919–36.

80. OMEGA version 2.4.3. OpenEye Scientific Software SF, NM. http://www.eyesopen.com.

81. Rush TS, 3rd, Grant JA, Mosyak L, Nicholls A. A shape-based 3-D scaffold hopping method and its application to a bacterial protein-protein interaction. J Med Chem. 2005; 48:1489–95.

82. ROCS version 3.2.0.3. OpenEye Scientific Software, Santa Fe, NM. http://www.eyesopen.com.

83. Irwin JJ, Sterling T, Mysinger MM, Bolstad ES, Coleman RG. ZINC: a free tool to discover chemistry for biology. J Chem Inf Model. 2012; 52:1757–68.

84. Grygorenko OO, Radchenko DS, Dziuba I, Chuprina A, Gubina KE, Moroz YS. Generating Multibillion Chemical Space of Readily Accessible Screening Compounds. iScience. 2020; 23:101681.

85. Baell J, Walters MA. Chemistry: Chemical con artists foil drug discovery. Nature. 2014; 513:481–3.

86. Kharas MG, Lengner CJ, Al-Shahrour F, Bullinger L, Ball B, Zaidi S, Morgan K, Tam W, Paktinat M, Okabe R, Gozo M, Einhorn W, Lane SW, Scholl C, Frohling S, Fleming M, Ebert BL, Gilliland DG, Jaenisch R, Daley GQ. Musashi-2 regulates normal hematopoiesis and promotes aggressive myeloid leukemia. Nat Med. 2010; 16:903–8.

87. Ito T, Kwon HY, Zimdahl B, Congdon KL, Blum J, Lento WE, Zhao C, Lagoo A, Gerrard G, Foroni L, Goldman J, Goh H, Kim SH, Kim DW, Chuah C, Oehler VG, Radich JP, Jordan CT, Reya T. Regulation of myeloid leukaemia by the cell-fate determinant Musashi. Nature. 2010; 466:765–8.

88. Li N, Yousefi M, Nakauka-Ddamba A, Li F, Vandivier L, Parada K, Woo DH, Wang S, Naqvi AS, Rao S, Tobias J, Cedeno RJ, Minuesa G, Y K, Barlowe TS, Valvezan A, Shankar S, Deering RP, Klein PS, Jensen ST, Kharas MG, Gregory BD, Yu Z, Lengner CJ. The Msi Family of RNA-Binding Proteins Function Redundantly as Intestinal Oncoproteins. Cell Rep. 2015; 13:2440–55.

89. Cimmperman P, Baranauskienė L, Jachimovičiūtė S, Jachno J, Torresan J, Michailovienė V, Matulienė J, Sereikaitė J, Bumelis V, Matulis D. A quantitative model of thermal stabilization and destabilization of proteins by ligands. Biophysical journal. 2008; 95:3222–31.

90. Layton CJ, Hellinga HW. Thermodynamic analysis of ligand-induced changes in protein thermal unfolding applied to high-throughput determination of ligand affinities with extrinsic fluorescent dyes. Biochemistry. 2010; 49:10831–41.

91. Morgan CE, Meagher JL, Levengood JD, Delproposto J, Rollins C, Stuckey JA, Tolbert BS. The First Crystal Structure of the UP1 Domain of hnRNP A1 Bound to RNA Reveals a New Look for an Old RNA Binding Protein. J Mol Biol. 2015; 427:3241–57.

92. Montemayor EJ, Curran EC, Liao HH, Andrews KL, Treba CN, Butcher SE, Brow DA. Core structure of the U6 small nuclear ribonucleoprotein at 1.7-A resolution. Nat Struct Mol Biol. 2014; 21:544–51.

93. Eibl JK, Strasser BC, Ross GM. Identification of novel pyrazoloquinazolinecarboxilate analogues to inhibit nerve growth factor in vitro. Eur J Pharmacol. 2013; 708:30–7.

94. Chakravarthy R, Mnich K, Gorman AM. Nerve growth factor (NGF)-mediated regulation of p75(NTR) expression contributes to chemotherapeutic resistance in triple negative breast cancer cells. Biochem Biophys Res Commun. 2016; 478:1541–7.

95. Erazo T, Evans CM, Zakheim D, Chu KL, Refermat AY, Asgari Z, Yang X, Da Silva Ferreira M, Mehta S, Russo MV, Knezevic A, Zhang XP, Chen Z, Fennell M, Garippa R, Seshan V, de Stanchina E, Barbash O, Batlevi CL, Leslie CS, Melnick AM, Younes A, Kharas MG. TP53 mutations and RNA-binding protein MUSASHI-2 drive resistance to PRMT5-targeted therapy in B-cell lymphoma. Nat Commun. 2022; 13:5676.

96. Karmakar S, Ramirez O, Paul KV, Gupta AK, Kumari V, Botti V, de Los Mozos IR, Neuenkirchen N, Ross RJ, Karanicolas J, Neugebauer KM, Pillai MM. Integrative genome-wide analysis reveals EIF3A as a key downstream regulator of translational repressor protein Musashi 2 (MSI2). NAR Cancer. 2022; 4:zcac015.

97. Yoon JH, Gorospe M. Identification of mRNA-Interacting Factors by MS2-TRAP (MS2-Tagged RNA Affinity Purification). Methods Mol Biol. 2016; 1421:15–22.

98. Park SM, Gonen M, Vu L, Minuesa G, Tivnan P, Barlowe TS, Taggart J, Lu Y, Deering RP, Hacohen N, Figueroa ME, Paietta E, Fernandez HF, Tallman MS, Melnick A, Levine R, Leslie C, Lengner CJ, Kharas MG. Musashi2 sustains the mixed-lineage leukemia-driven stem cell regulatory program. J Clin Invest. 2015; 125:1286–98.

99. Vu LP, Prieto C, Amin EM, Chhangawala S, Krivtsov A, Calvo-Vidal MN, Chou T, Chow A, Minuesa G, Park SM, Barlowe TS, Taggart J, Tivnan P, Deering RP, Chu LP, Kwon JA, Meydan C, Perales-Paton J, Arshi A, Gonen M, Famulare C, Patel M, Paietta E, Tallman MS, Lu Y, Glass J, Garret-Bakelman FE, Melnick A, Levine R, Al-Shahrour F, Jaras M, Hacohen N, Hwang A, Garippa R, Lengner CJ, Armstrong SA, Cerchietti L, Cowley GS, Root D, Doench J, Leslie C, Ebert BL, Kharas MG. Functional screen of MSI2 interactors identifies an essential role for SYNCRIP in myeloid leukemia stem cells. Nat Genet. 2017; 49:866–75.

100. Zhang H, Tan S, Wang J, Chen S, Quan J, Xian J, Zhang S, He J, Zhang L. Musashi2 modulates K562 leukemic cell proliferation and apoptosis involving the MAPK pathway. Exp Cell Res. 2014; 320:119–27.

101. Kudinov AE, Deneka A, Nikonova AS, Beck TN, Ahn YH, Liu X, Martinez CF, Schultz FA, Reynolds S, Yang DH, Cai KQ, Yaghmour KM, Baker KA, Egleston BL, Nicolas E, Chikwem A, Andrianov G, Singh S, Borghaei H, Serebriiskii IG, Gibbons DL, Kurie JM, Golemis EA, Boumber Y. Musashi-2 (MSI2) supports TGF-beta signaling and inhibits claudins to promote non-small cell lung cancer (NSCLC) metastasis. Proc Natl Acad Sci U S A. 2016; 113:6955–60.

102. Jafari R, Almqvist H, Axelsson H, Ignatushchenko M, Lundback T, Nordlund P, Martinez Molina D. The cellular thermal shift assay for evaluating drug target interactions in cells. Nat Protoc. 2014; 9: 2100–22.

103. Nissen P, Kjeldgaard M, Nyborg J. Macromolecular mimicry. EMBO J. 2000; 19:489–95.

104. Tsonis PA, Dwivedi B. Molecular mimicry: structural camouflage of proteins and nucleic acids. Biochim Biophys Acta. 2008; 1783:177–87.

105. Della Volpe S, Nasti R, Queirolo M, Unver MY, Jumde VK, Dömling A, Vasile F, Potenza D, Ambrosio FA, Costa G. Novel Compounds Targeting the RNA-Binding Protein HuR. Structure-Based Design, Synthesis, and Interaction Studies. ACS medicinal chemistry letters. 2019; 10:615–20.

106. Baek M, McHugh R, Anishchenko I, Baker D, DiMaio F. Accurate prediction of nucleic acid and protein-nucleic acid complexes using RoseTTAFoldNA. bioRxiv. 2022.

107. Sakamoto KM, Kim KB, Kumagai A, Mercurio F, Crews CM, Deshaies RJ. Protacs: chimeric molecules that target proteins to the Skp1-Cullin-F box complex for ubiquitination and degradation. Proc Natl Acad Sci U S A. 2001; 98:8554–9.

108. Pettersson M, Crews CM. PROteolysis TArgeting Chimeras (PROTACs)—past, present and future. Drug Discovery Today: Technologies. 2019.

