## Supporting Information for "Rationally designed inhibitors of the Musashi protein-RNA interaction by hotspot mimicry"

[†]Equal author contributions.

### Supporting Methods

*PDB structures used in calculations*

The calculations that led to selection of R1-R12 were carried out using model 1 of the NMR structure of Musashi-1 bound to RNA (PDB ID 2RS2) [S1].

*Building hotspot pharmacophores*

Hotspot pharmacophores were built using a new dedicated protocol implemented in the Rosetta software suite [S2], and is freely available for academic use (www.rosettacommons.org). The hotspot pharmacophore is extracted solely from the structure of the protein-RNA complex, and thus does not take into any account potential rearrangement of the protein upon RNA binding.

To select deeply buried RNA bases, the solvent accessible surface area (SASA) of each base in the RNA was calculated in the presence of the protein, then re-calculated after deleting the protein: this yielded the SASA that was directly buried by the protein in the complex. A base was carried forward if the change in SASA upon complexation was greater than a preset cutoff value (46.81 $Å^2$ for adenine, 31.09 $Å^2$ for cytosine, 45.06 $Å^2$ for guanine and 52.66 $Å^2$ for uracil); these values correspond to the median values of 344 non-redundant protein-RNA complexes retrieved from the Protein-RNA Interface Database (PRIDB) [S3] in March 2013 (http://pridb.gdcb.iastate.edu/download/RB344.txt).

Polar groups from the RNA that participate in intermolecular hydrogen bonding (as defined using the Rosetta energy function) are also included.

The resulting interaction maps are then clustered using a modified version of Kruskal's minimum spanning tree algorithm. We first build a complete graph, in which vertices are the ring moieties, and the edge weights are the Euclidean distances between vertices. Then we take edges in ascending order and cluster the end vertices of that edge if no cycle would be caused. We halt the clustering when the distance is greater than a user-specified cutoff value (default 5.0 Å). The donor/acceptor atoms are then assigned to the closest ring moieties if the distance is less than another user-specified value (default 5.0 Å). Finally, we output the pharmacophore templates if the cluster contains at least two ring moieties.

The Rosetta command line used to carry out this new functionality is as follows:

```
gen_rna_pharmacophore.macosgccrelease -input_protein xxx_protein.pdb -input_rna xxx_rna.pdb
```

*Identifying complementary ligands*

We used ROCS to screen large libraries for compounds that match the hotspot pharmacophore. We downloaded the standard 'drugs-now' subset of ~7 million molecules from ZINC database for screening [S4]. We generated up to 100 conformers for each molecule in the database using OMEGA [S5-7]. We screened the database using the hotspot pharmacophore (using default ROCS parameters), and carried forward the top 500 compounds ranked by 'TanimotoCombo' score. We then aligned these back to the protein using the hotspot pharmacophore, then carried out a gradient-based full-atom minimization of the complex using the Rosetta energy function [S2]. This energy function includes terms that capture packing, hydrogen bonding, implicit solvation (modeled via EEF1 [S8]), sidechain rotamer preferences, and backbone dihedral preferences. After minimization, the top-scoring compounds were visually inspected and selected for experimental validation based on cost and availability.

*Predicting target selectivity*

The complete set of 1792 protein-RNA complexes was retrieved from the PDB in June 2014. Hotspot pharmacophores were extracted from each complex, and non-unique pharmacophores were removed (those with ROCS shape_tanimoto > 0.94 and color_tanimoto > 0.74). This left 543 unique pharmacophores that were comprised of at least two rings, derived from 362 different protein-RNA complexes.

Conformers for each compound were generated by OMEGA using the following command line:

```
omega2 -in xxx.pdb -strictatomtyping false -strictstereo false -strictfrags false -searchff mmff94s -buildff mmff94s -maxconfs 500
```

For a given compound, we then used ROCS to screen conformers of this molecule against the library of hotspot pharmacophores using the following command line:

```
rocs -dbase conformer_ensemble.pdb -query hotspot.pdb -oformat pdb -rankby FitTverskyCombo
```

The multidimensional scaling (MDS) analysis presented in Figure 5 was carried out in R [S9], using the "cmdscale" function. Pharmacophores for hnRNP A1 and Prp24 were extracted from PDB IDs 4YOE and 4N0T, respectively.

*Compound optimization*

We carried out two sequential rounds of optimization via automated fingerprint-based searches, followed by a round of traditional medicinal chemistry optimization by combining desirable fragments and R-groups from the best compounds from earlier rounds of optimization. For each of the rounds of the fingerprint-based optimization, we took advantage of Enamine Real Database, a database of greater than 11 billion compounds (at the time) that are readily synthesizable and sold at affordable pricing. We started by querying the database with our initial hit molecule from screening ZINC library (R12), based on fingerprint similarity score, we selected the top 1000 compounds. From these, we clustered based on diversity of substituent on the rings and linker length and select representative compounds of the clusters for biochemical characterization. In the first round of screening, we choose 16 compounds; in the second round, we selected 50 diverse compounds based on similarity to the best compound from the first round. Finally, the knowledge acquired from the structure activity relationship of the 66 compounds tested so far was combined in designing new inhibitors; based on these, we purchased 7 additional compounds for biochemical characterization.

*Model building of R12 derivatives*

For each round of optimization carried out, we built structural models for the compounds to assess whether the compounds are making appropriate interaction with the protein. For this, we generated 300 low energy 3D conformations for each of the compounds using OpenEye OMEGA[S5-7]. Then, we aligned each of the conformers to Msi1 pharmacophore (generated from model 1 of the NMR structure with PDB ID 2RS2) using OpenEye ROCS[S10]; then, for each compound, we selected top 10 conformers with the highest shape and electrostatic overlap with the pharmacophore (TanimotoCombo). We placed these in the binding site of the protein and energy-minimized the complexes using Rosetta energy function. Finally, for each compound, we selected the conformer with the lowest energy.

*Protein expression and purification*

The RRM1-RRM2 domain of human Msi1 and Msi2 were purchased from Genewiz as a fusion protein with an N-terminal 6xHis-tag and a tobacco etch virus (TEV) protease site on vector pET28a (+). The RRM1 domain of human Msi1 with an N-terminal 6xHis-tagged streptococcal GB1 domain and hnRNP A1 with an N-terminal 6xHis-tagged maltose- binding protein (MBP) fusion proteins were purchased from Genewiz, both on vector pET28a (+). Each of these four constructs were expressed and purified as described below.

The expression plasmid was transformed into *Escherichia coli* BL21(DE3) pLysS, then a 5 mL overnight starter culture was used to inoculate a 1 L culture of Luria-Bertani (LB) media. Cells were grown at 37 ºC to an $OD_{600}$ of 0.6–0.8 and were induced with 1 mM IPTG at 37 ºC for 4 hours. The induced cells were harvest and the pellet was resuspended in lysis buffer (20 mM HEPES, 1 M NaCl, 50 mM imidazole, 1 mM DTT, pH 7.4) and sonicated for 10 minutes (Fisher Scientific Sonic Dismembrator Model 100). The cell lysates were then centrifuged at 15,000g for 50 min. The protein of interest remained in the supernatant, which was purified by HPLC affinity chromatography with Ni-chelated Sepharose Fast Flow Resin (GE Healthcare). The buffer was exchanged with dialysis (20 mM HEPES, 150 mM NaCl, 0.1 mM EDTA, 1 mM DTT, pH 7.4).

All protein concentrations were determined with reference to bovine albumin standards using Bradford assays.

*Fluorescence polarization competition assays*

A fluorescently labeled RNA oligonucleotide corresponding to the *NUMB* sequence (UAGGUAGUAGU/36-FAM/) was purchased from Integrated DNA Technologies (Coralville, IA) and dissolved in RNase free water. To measure the dissociation constant of Msi1 RRM1-RRM2 and RNA binding, a fixed concentration (5 nM) of fluorescein-labeled RNA and increasing concentrations of Msi1 RRM1-RRM2 (0 nM to 128 nM) were mixed in binding assay buffer (20 mM HEPES, 150 mM NaCl, 0.01% Triton-X100 pH 7.4). Fluorescence intensities were measured in replicate on the Molecular Devices SpectraMax® i3x (San Jose, CA) and the fluorescence polarization value (FP) was calculated by the following equation:

$$FP = \frac{I_{\parallel} - I_{\perp}}{I_{\parallel} + I_{\perp}}$$

where $I_{\parallel}$ refers to the intensity of the parallel fluorescence and $I_{\perp}$ refers to the intensity of the perpendicular fluorescence. The dissociation constant ($K_D$) was fit using Prism 6 (GraphPad Software Inc.) as follows:

$$Y = Bottom + \frac{Top - Bottom}{1 + EC_{50}^{Hill\ Slope} / L^{Hill\ Slope}}$$

To examine the displacement of RNA by R12 derivative compounds, the competition assays were performed with 8 nM as fixed Msi1 concentration, 5 nM as fixed fluorescein-labeled RNA concentration, and a serial dilution of compounds were added. Data were fit to a single-site competition model to determine IC50 using Prism 6 (GraphPad Software Inc.). There are two RRM domains in the protein construct therefore the Hill coefficient was set to be free. Given the known experimental conditions and the binding affinity described above, the Ki was then computed from the IC50 using the method of Nikolovska-Coleska et al [S11].

To test the contribution to binding affinity of each base in the *NUMB* RNA sequence, we purchased eight shorter oligos that one with the fluorescein-label, five of them harbor an abasic site at a different position, as well as the corresponding wild-type (**Table S2**). The shorter fluorescein-labeled RNA was applied here to match the other RNA fragments with the abasic site. $K_i$ values were then determined from the competition experiment described as above.

*Differential scanning fluorimetry (Thermofluor)*

Differential scanning fluorimetry (DSF) experiments were carried out using a standard protocol described by others [S12]. The protein concentration was fixed to be 4 μM and SYPRO Orange (Invitrogen S6651) was used at a final concentration of 5X. The experiments were carried out in 20 mM HEPES, 150 mM NaCl, 0.01% Triton-X100, 2.5% DMSO pH 7.4, with Eppendorf Realplex2 Mastercycler. Each sample was divided to three 50 μL replicates. Sample solutions were dispensed into 96-well optical reaction plate (Thermo Fisher Scientific 4306737) and the plate was sealed with optical PCR plate sheet (Thermo Fisher Scientific AB-1170). Fluorescence intensity was measured via the JOE emission filter (550 nm) and "PTS clear plate" was set as the background for the calibration. Temperature was continuously increased: 0.4 ºC/min, from 37 ºC to 56.6 ºC. Melting curves were directly exported from the instrument, and then were analyzed with Prism 6 (GraphPad Software Inc.).

*Nuclear magnetic resonance (NMR) spectroscopy*

$^{15}$N-labeled protein was expressed and purified as described above then cleaved with TEV overnight at 4 ºC in 20 mM HEPES pH 6.3, 50 mM NaCl, and 2 mM DTT in a 1:20 ratio. Cleaved protein was then passed over a 5 mL HisTrap column. Pure fractions were then pooled and concentrated to 1 mL. Buffer exchange was performed using a NAP10 column (GE Healthcare) into 20mM HEPES pH 7.0, 1 mM TCEP and 10% $D_2O$.

All spectra were recorded at 298K on a Bruker Ascend 600-MHz spectrometer. DMSO control spectra were prepared by adding DMSO to a final concentration of 0.5% in a 75 μM protein solution. Compound R12.8.44.3 were added to a final concentrations of 100 μM, respectively. All data were processed using TopSpin 4.0.

*Aptamer pull-down reporter assay*

HEK293T cells were seeded at a density of 300,000 cells per ml in a 10 cm tissue culture dish 24 hour prior to transfection. Transient transfection with pMS2-LUC-3'UTR [S13] was performed using the TransIT-X2 delivery system (Mirus). Cells were incubated for 2 hours following transfection at 37 ºC in a 5% $CO_2$ humidified chamber, for uptake of plasmid. After this incubation, the compound of interest (or equivalent volume of DMSO) was added. After 24 hours, cells were harvested. Immunoprecipitation was carried out using Anti-FLAG M2 affinity gel (Sigma) as previously described [S13]. Bound complexes were eluted using 0.4 ug/ml of 3x FLAG peptide. Input and IP samples were denatured in NuPAGE LDS sample buffer (Invitrogen) for 10 min in boiling water, then resolved on a 4-20% SDS page gel followed by wet transfer onto 0.2 μM nitrocellulose membrane (Biorad) for 90 min at 100 V at 4 ºC. The membrane was blocked in 5% nonfat milk/TBS-T and probed with the antibodies as indicated. The membrane was developed with highly sensitive Radiance Plus chemiluminescent substrate (Azure Biosystems).

*Cell culture*

PC9 cells (human alveolar basal epithelial adenocarcinoma cell line bearing EGFR exon 19 deletion) were obtained from the American Type Culture Collection (ATCC). K562 cells (human chronic myelogenous leukemia line with TP53 (p.Q136fs*13) and CDKN2A deletion) were obtained from the Leonidas Platanias group (Northwestern University Lurie Cancer Center). Initial stocks were cryopreserved, and at every 6-month interval a fresh aliquot of frozen cells was used for experiments. All cells were cultured in RPMI 1640 (Gibco, Gaithersburg, MD) supplemented with 10% FBS (Hyclone, Logan, UT), penicillin (100U/ml), streptomycin (100μg/ml), sodium pyruvate (1 mM) and non-essential amino acids (0.1 mM) under conditions indicated in the figure legends.

Ro 08-2750 was obtained from Tocris (#2272, Minneapolis, MN), and osimertinib was obtained from SelleckChem (#S7297, Houston, TX).

*Western blot analysis*

Preparation of cell lysates and Western blot analysis were performed using standard methods as previously described [S14]. Bands signals were detected by X-ray films, and films were digitized by photo scanner. Image analysis was done using ImageJ (version 1.53e, National Institutes of Health, Bethesda, MD), with signal intensity normalized to β-actin. 3 or 4 replicates were used for quantitation of each experiment.

Anti-Msi2 (#ab76148) was obtained from Abcam (Cambridge, UK). Anti-EGF receptor (#4267), anti-β-actin (#3700), anti-SMAD3 (#9513), anti-rabbit HRP-linked (#7074), anti-mouse HRP-linked (#7076) were obtained from Cell Signaling (Danvers, MA).

*Cellular thermal shift assay (CETSA)*

Cells were incubated with the compound of interest for 3 hours in complete media. Cells were harvested and washed in PBS with protease inhibitor cocktail (cat#78430, ThermoFisher Scientific). Cell pellets were heated for 3 minutes at the desired temperature in a BioRad 96-well thermal cycler (cat#1861096), then immediately the tubes were removed and incubated at room temperature for 3 min. Cells were then lysed by three rounds of freeze-thawing (alternating exposure of the samples to liquid nitrogen and 20 °C in the thermal cycler). Before Western blotting, aggregated protein was removed by centrifugation at 20,000 g for 20 min at 4 °C and collecting the supernatant. The supernatant was then analyzed directly by Western blot [S15].

*RNA IP (RIP) assay*

RNA was immunoprecipitated from cell lysates ($2x10^7$ cells per IP) using either a control normal rabbit IgG or rabbit monoclonal anti-Msi2 antibody (#ab76148) and the Magna RIP RNA-binding Protein Immunoprecipitation kit (cat#17-700, Millipore, Burlington, MA) using manufacturer's protocol. Immunoprecipitated RNAs were quantified by qPCR using primers indicated (**Table S3**), using PTP4A1 as a normalization (positive) control and GAPDH as a negative control.

*Reverse transcription and qPCR*

RNA was extracted using phenol-chloroform based method. RNA concentration and quantity was measured using NanoDrop Lite (cat# ND-LITE ThermoFisher Scientific). First strand cDNA synthesis was performed with iScript cDNA synthesis kit (cat#1708841, Biorad, California, USA) according to manufacturer's instructions. The generated cDNA was diluted tenfold and used as a template for qPCR, which was performed with Applied Biosystems QuantStudio 3 system using PowerTrack™ SYBR Green Master Mix (Applied Biosystems). Relative quantification of genes expression was performed using $2^{-\Delta\Delta Ct}$ method.

*Cell viability assay*

To analyze the effect of compound treatment on cell proliferation, cells were plated (500 cells/well) in 96-well cell culture plates in complete media. We used increasing concentrations of compounds to calculate $IC_{50}$ values for each cell line. After 72 hours incubation with a given compound, we used CellTiter-Blue® assay (Promega, Fitchburg, WI) and measured OD for the supernatant at 562 nm. Data were analyzed using GraphPad Prism.

*Clonogenic assay*

PC9 cells (200 cells per well) were plated in 12-well pates and incubated in complete media. After 24 hours compounds were added to evaluate their effects on cell proliferation. Cells were incubated for 7 days. Cells were then fixed in 10% acetic acid/10% methanol solution and stained with 0.5% (w/v) crystal violet as previously described [S16]. A colony was defined as consisting of >50 cells and counted digitally using ImageJ software as described previously [S17].

### Supporting References


S1. Ohyama T, Nagata T, Tsuda K, Kobayashi N, Imai T, Okano H, Yamazaki T, Katahira M. Structure of Musashi1 in a complex with target RNA: the role of aromatic stacking interactions. Nucleic Acids Res. 2012;40(7):3218-31.

S2. Leaver-Fay A, Tyka M, Lewis SM, Lange OF, Thompson J, Jacak R, Kaufman K, Renfrew PD, Smith CA, Sheffler W, Davis IW, Cooper S, Treuille A, Mandell DJ, Richter F, Ban YE, Fleishman SJ, Corn JE, Kim DE, Lyskov S, Berrondo M, Mentzer S, Popovic Z, Havranek JJ, Karanicolas J, Das R, Meiler J, Kortemme T, Gray JJ, Kuhlman B, Baker D, Bradley P. ROSETTA3: an object-oriented software suite for the simulation and design of macromolecules. Methods Enzymol. 2011;487:545-74.

S3. Lewis BA, Walia RR, Terribilini M, Ferguson J, Zheng C, Honavar V, Dobbs D. PRIDB: a Protein-RNA interface database. Nucleic Acids Res. 2011;39(Database issue):D277-82.

S4. Irwin JJ, Sterling T, Mysinger MM, Bolstad ES, Coleman RG. ZINC: A Free Tool to Discover Chemistry for Biology. J Chem Inf Model. 2012.

S5. Hawkins PC, Skillman AG, Warren GL, Ellingson BA, Stahl MT. Conformer generation with OMEGA: algorithm and validation using high quality structures from the Protein Databank and Cambridge Structural Database. J Chem Inf Model. 2010;50(4):572-84.

S6. Hawkins PC, Nicholls A. Conformer generation with OMEGA: learning from the data set and the analysis of failures. J Chem Inf Model. 2012;52(11):2919-36.

S7. OMEGA version 2.4.3. OpenEye Scientific Software SF, NM. http://www.eyesopen.com.

S8. Lazaridis T, Karplus M. Effective energy function for proteins in solution. Proteins. 1999;35(2):133-52.

S9. R Core Team. R: A Language and Environment for Statistical Computing. Vienna, Austria: R Foundation for Statistical Computing; 2014.

S10. ROCS version 3.2.0.3. OpenEye Scientific Software, Santa Fe, NM. http://www.eyesopen.com.

S11. Nikolovska-Coleska Z, Wang R, Fang X, Pan H, Tomita Y, Li P, Roller PP, Krajewski K, Saito NG, Stuckey JA, Wang S. Development and optimization of a binding assay for the XIAP BIR3 domain using fluorescence polarization. Anal Biochem. 2004;332(2):261-73.

S12. Niesen FH, Berglund H, Vedadi M. The use of differential scanning fluorimetry to detect ligand interactions that promote protein stability. Nat Protoc. 2007;2(9):2212-21.

S13. Karmakar S, Ramirez O, Paul KV, Gupta AK, Kumari V, Botti V, de Los Mozos IR, Neuenkirchen N, Ross RJ, Karanicolas J, Neugebauer KM, Pillai MM. Integrative genome-wide analysis reveals EIF3A as a key downstream regulator of translational repressor protein Musashi 2 (MSI2). NAR Cancer. 2022;4(2):zcac015.

S14. Kudinov AE, Deneka A, Nikonova AS, Beck TN, Ahn YH, Liu X, Martinez CF, Schultz FA, Reynolds S, Yang DH, Cai KQ, Yaghmour KM, Baker KA, Egleston BL, Nicolas E, Chikwem A, Andrianov G, Singh S, Borghaei H, Serebriiskii IG, Gibbons DL, Kurie JM, Golemis EA, Boumber Y.

Musashi-2 (MSI2) supports TGF-beta signaling and inhibits claudins to promote non-small cell lung cancer (NSCLC) metastasis. Proc Natl Acad Sci U S A. 2016;113(25):6955-60.

S15. Jafari R, Almqvist H, Axelsson H, Ignatushchenko M, Lundback T, Nordlund P, Martinez Molina D. The cellular thermal shift assay for evaluating drug target interactions in cells. Nat Protoc. 2014;9(9):2100-22.

S16. Abazeed ME, Adams DJ, Hurov KE, Tamayo P, Creighton CJ, Sonkin D, Giacomelli AO, Du C, Fries DF, Wong KK, Mesirov JP, Loeffler JS, Schreiber SL, Hammerman PS, Meyerson M. Integrative radiogenomic profiling of squamous cell lung cancer. Cancer Res. 2013;73(20):6289-98.

S17. Cai Z, Chattopadhyay N, Liu WJ, Chan C, Pignol JP, Reilly RM. Optimized digital counting colonies of clonogenic assays using ImageJ software and customized macros: comparison with manual counting. Int J Radiat Biol. 2011;87(11):1135-46.

S18. Robert X, Gouet P. Deciphering key features in protein structures with the new ENDscript server. Nucleic Acids Res. 2014;42(Web Server issue):W320-4.

S19. ESPript - http://espript.ibcp.fr.

### Supporting Tables

| Compound | 2D Structure | EC50 (μM) in Fluorescence Polarization Assay |
|---|---|---|
| **R12** | 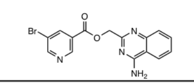 | N.D.* |
| **R12-7** | 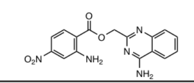 | 49 |
| **R12-8** | 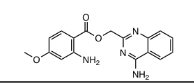 | > 100 |
| **R12-8-19** | 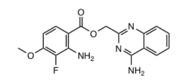 | > 100 |
| **R12-8-22** | 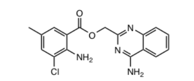 | 90 |
| **R12-8-29** | 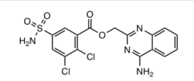 | 100 |
| **R12-8-38** | 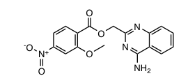 | > 100 |
| **R12-8-44** | 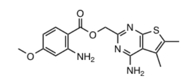 | 22 |
| **R12-8-46** | 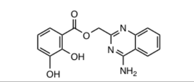 | 9 |
| **R12-8-47** | 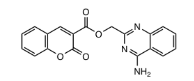 | > 100 |
| **R12-8-48** | 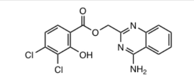 | > 100 |
| **R12-8-44-1** | 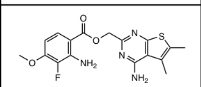 | 16 |
| **R12-8-44-2** | 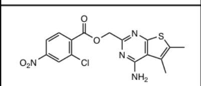 | > 100 |
| **R12-8-44-3** | 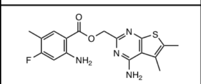 | 9 |
| **R12-8-44-4** | 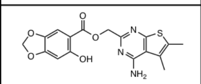 | 6 |
| **R12-8-44-6** | 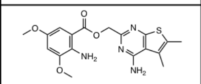 | > 100 |
| **R12-8-44-7b** | 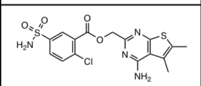 | > 100 |
| **R12-8-44-lk2** | 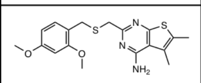 | 26 |

**Table S1: Three rounds of compound optimization, starting from initial hit R12.**

| Name | Sequence |
|---|---|
| FC-NUMB | 5'- F-GUAGU -3' |
| NUMBa0 (WT) | 5'- UGUAGUU -3' |
| NUMBa1 (G104x) | 5'- UxUAGUU -3' |
| NUMBa2 (U105x) | 5'- UGxAGUU -3' |
| NUMBa3 (A106x) | 5'- UGUxGUU -3' |
| NUMBa4 (G107x) | 5'- UGUAxUU -3' |
| NUMBa5 (U108x) | 5'- UGUAGxU -3' |

**Table S2: Sequences of RNA oligonucleotides used in biochemical assays.** "F" refers to the fluorescein label, and "x" refers to an abasic site (i.e. internal RNA spacer site).

| Gene Symbol | SYBR Green primers |
|---|---|
| *PTP4A* | Fw: 5`-ATCCAACCAATGCGACCTTA<br>Rev: 5`-AAGGCCAATCAAGAACATGG |
| *GAPDH* | Fw: 5`-TGCACCACCAACTGCTTAGC<br>Rev: 5`-GGCATGGACTGTGGTCATGAG |
| *EGFR* | Fw: 5`- CCCGTAATTATGTGGTGACAGA<br>Rev: 5`- ACCAATACCTATTCCGTTACACA |
| *SMAD3* | Fw: 5`-CCAGCACATAATAACTTGGACCT<br>Rev: 5`-GATGTGTCTCCGTGTCAGCTC |
| *MSI2* | Fw: 5`-GGTCATGAGAGATCCCACTACG<br>Rev: 5`-TCTACACTTGCTGGGTCTGC |

**Table S3: Primers used in cellular studies.** List of primers used to quantify gene expression in RT-PCR (SYBR Green) assays and in RNA immunoprecipitation assays.

### Supporting Figures

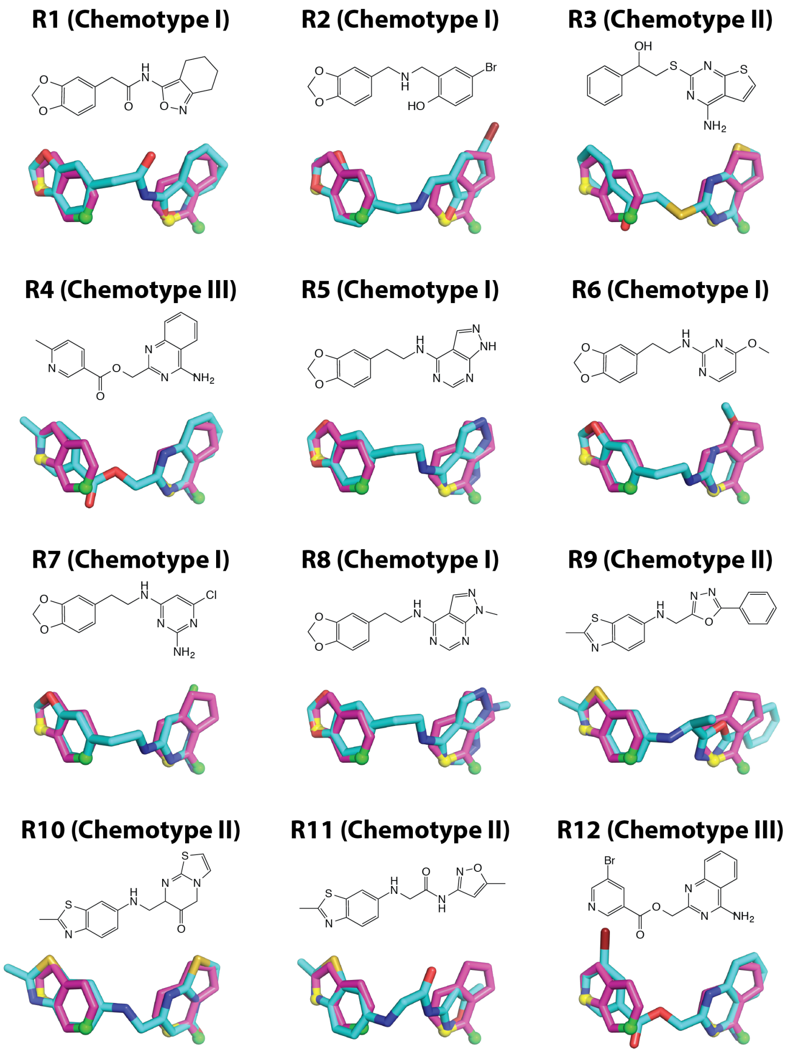


**Figure S1: The 12 compounds selected from initial computational screening.** The chemical structure is shown for each compound, as well as a three-dimensional model of each compound (*cyan*) superposed with the Msi1 RBD1 hotspot pharmacophore (*magenta*).

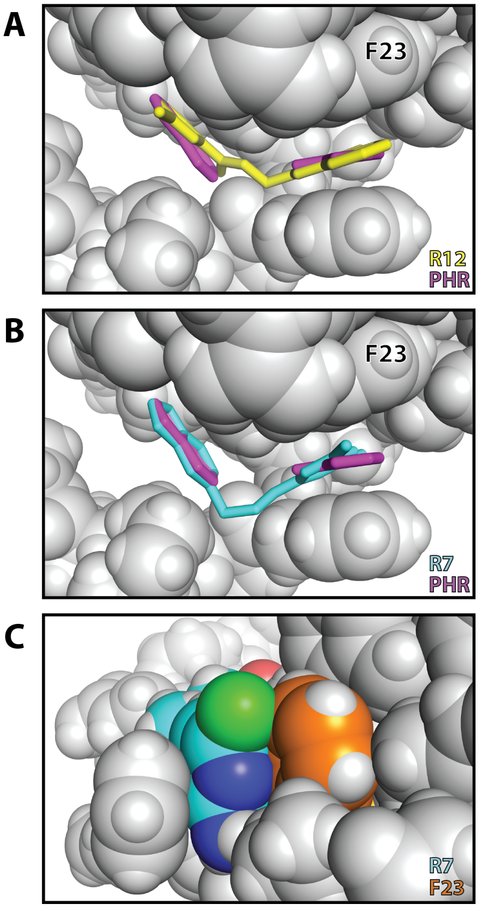


**Figure S2: An inadvertent steric clash may explain the lack of binding by R7. (A)** The rings in the model of R12 (*yellow*) are well-superposed with those of the hotspot pharmacophore (*magenta*), allowing for aromatic stacking with Msi1. **(B)** The relative positioning of the rings in the R7 (*cyan*) do not quite align with the hotspot pharmacophore (*right side of this perspective*). **(C)** This difference in the positioning of the ring leads to a steric clash with Phe23 (*orange*).

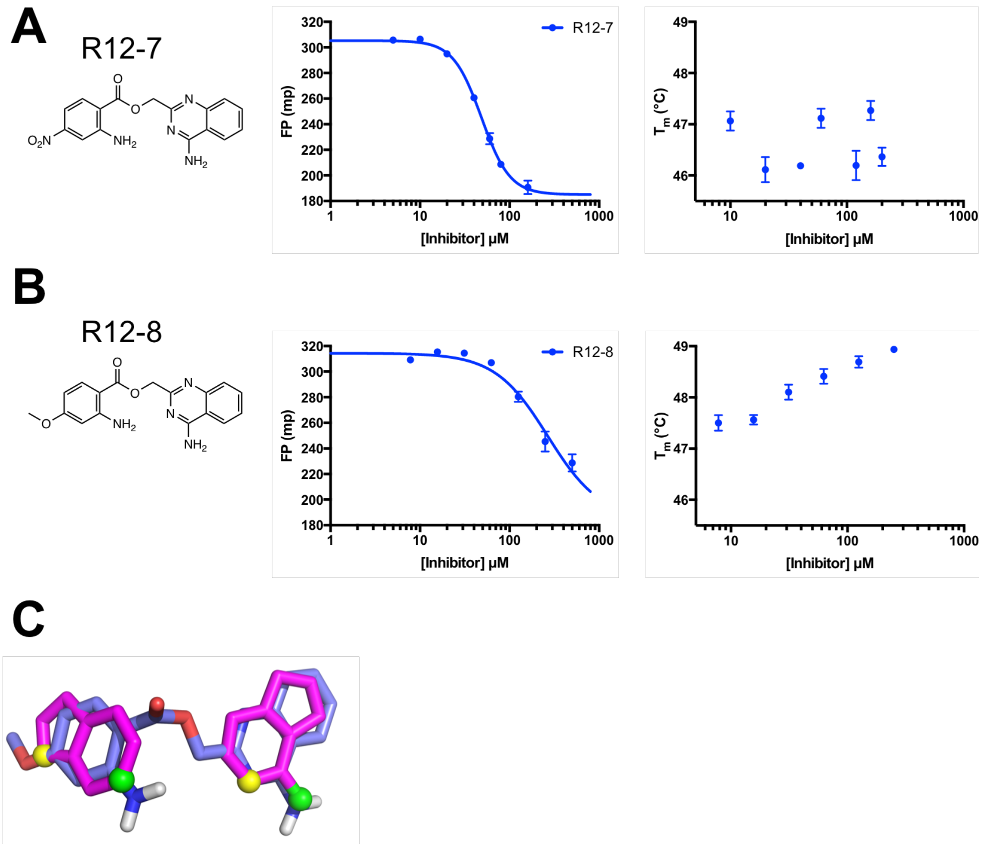


**Figure S3: Inhibitors coming from the first round SAR study. (A)** The chemical structure of R12-7 and the results of biochemical assays. The middle panel is the FP competition assay and the right panel is the DSF assay. **(B)** The chemical structure of R12-8 and the results of biochemical assays. The middle panel is the FP competition assay and the right panel is the DSF assay. **(C)** Superposition of R12-8 (*slate*) and the hotspot pharmacophore (*magenta*).

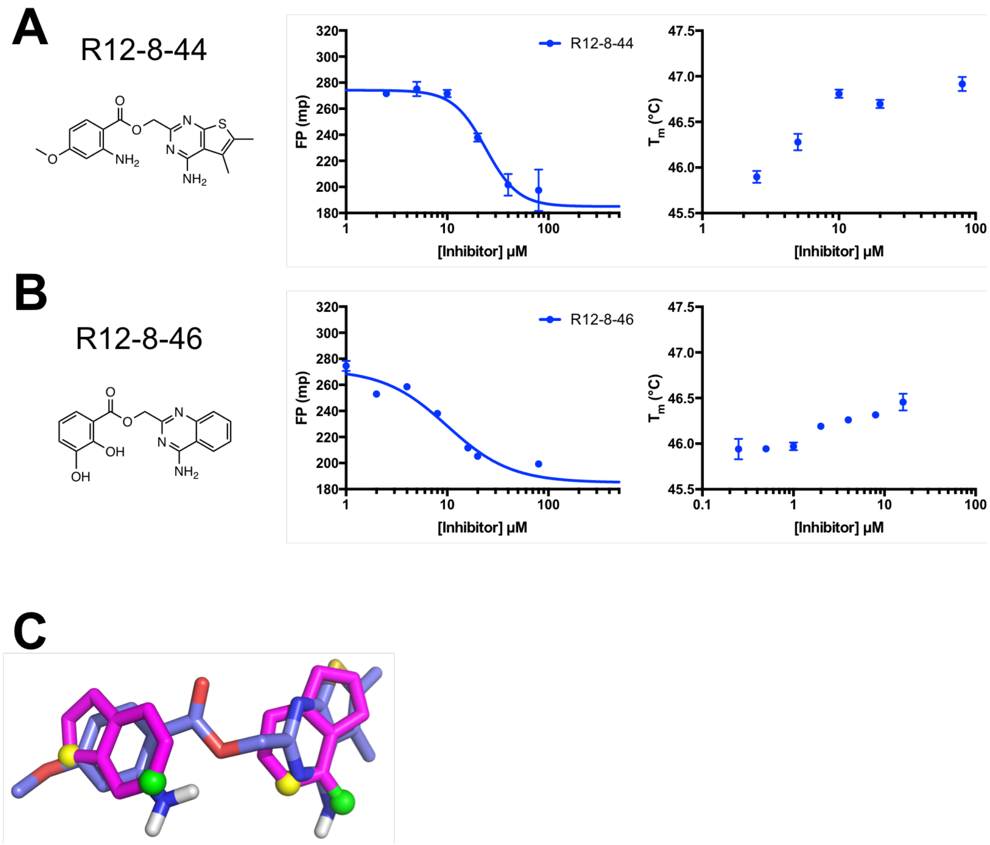


**Figure S4: Inhibitors coming from the second round SAR study. (A)** The chemical structure of R12-8-44 and the results of biochemical assays. The middle panel is the FP competition assay and the right panel is the DSF assay. **(B)** The chemical structure of R12-8-46 and the results of biochemical assays. The middle panel is the FP competition assay and the right panel is the DSF assay. **(C)** Superposition of R12-8-44 (*slate*) and the hotspot pharmacophore (*magenta*).

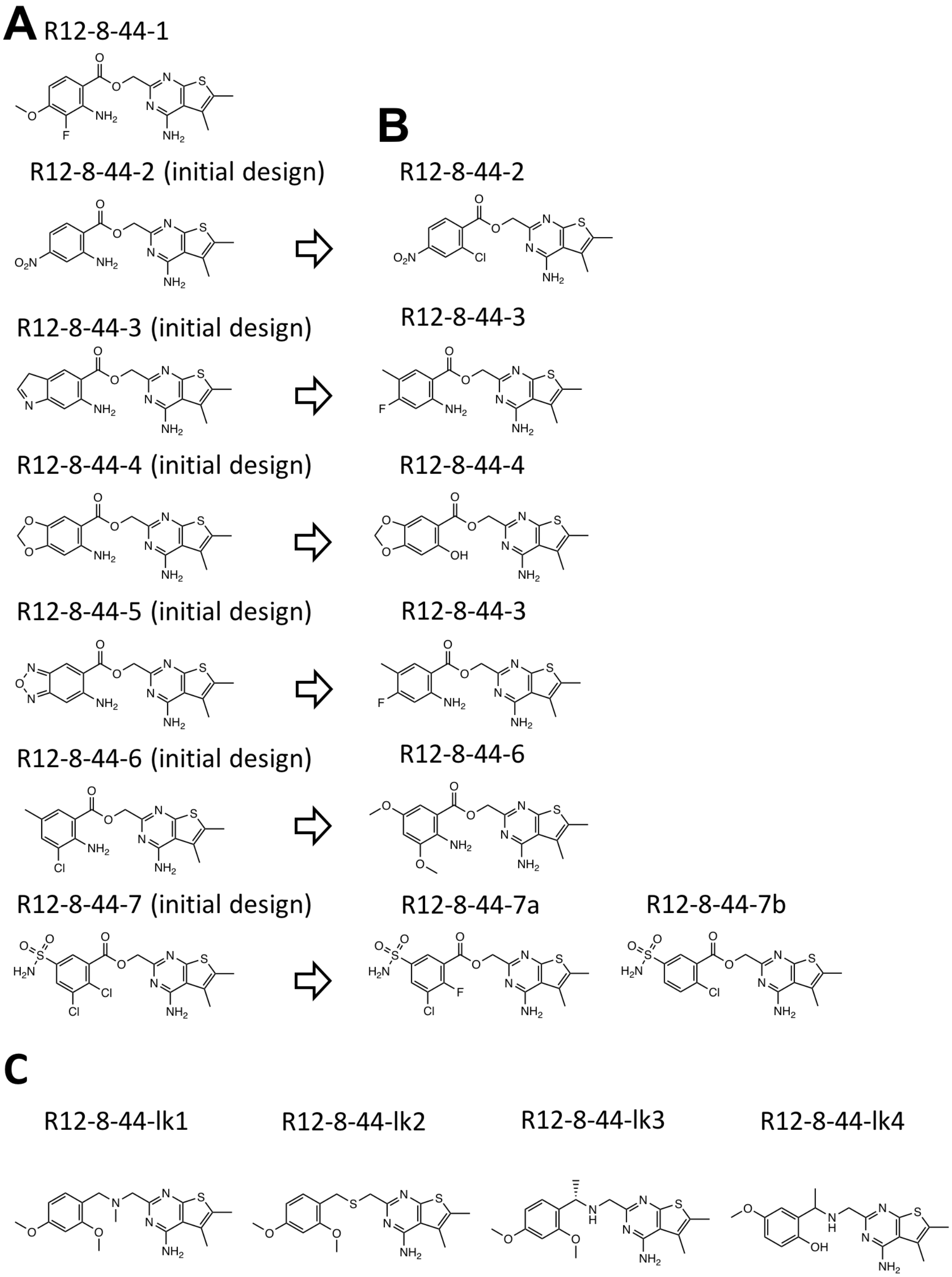


**Figure S5: Compounds design of the third round SAR study. (A)** Chemical structures of the initial compounds designed based on R12-8-44. **(B)** Chemical structures of the compounds which were commercially available. **(C)** Chemical structures of the compounds with similar structures as R12-8-44 but different linkers.

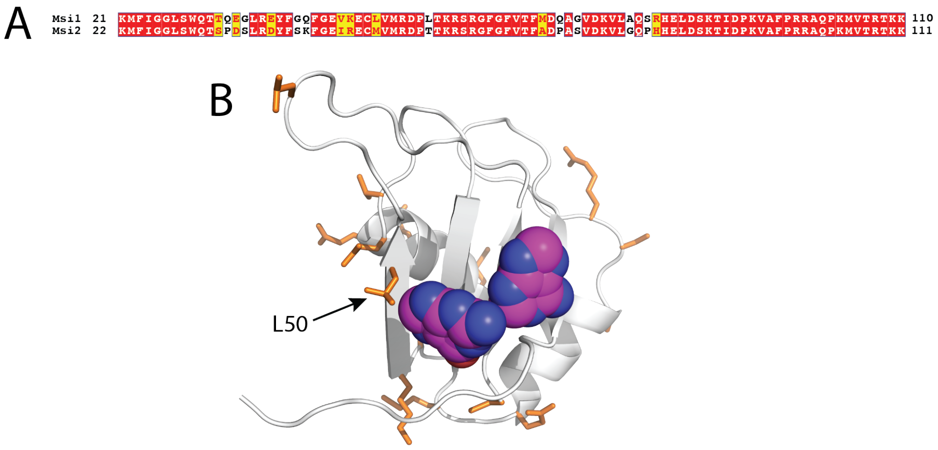


**Figure S6: Comparison of Msi1 and Msi2. (A)** Sequence alignment comparing RRM1 from Msi1 to RRM1 from Msi2. Over these 90 residues, only 17 positions differ (81% sequence identity) and only 9 correspond to non-conservative mutations (90% sequence similarity). This alignment was rendered using ESPript [S18,19]. **(B)** The structure of Msi1 RRM1 is shown (*grey cartoons*), with the hotspot pharmacophore derived from its cognate RNA (*magenta and blue spheres*). Residues at which the sequence differs in Msi2 RRM1 are highlighted (*orange sticks*); with the exception of Leu50 (Met in Msi2), each of the residues that differ are surface exposed and located far from the hotspot pharmacophore.

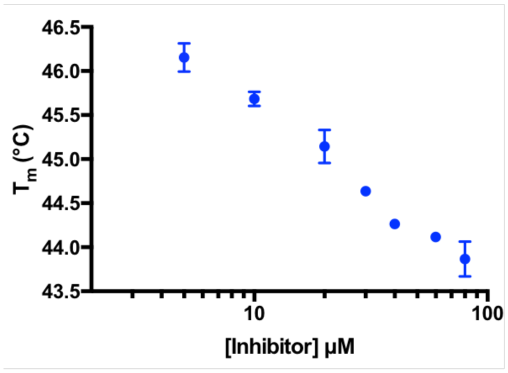


**Figure S7: DSF characterization of R12-8-44-3.** Differential scanning fluorimetry shows that addition of R12-8-44-3 decreases the melting temperature of Msi1 in a concentration-dependent manner. In light of NMR HSQC characterization showing that specific peaks respond to addition of this compound, we surmise that the unexpected response in this DSF experiment is due to a partially-folded intermediate in the thermal unfolding transition.

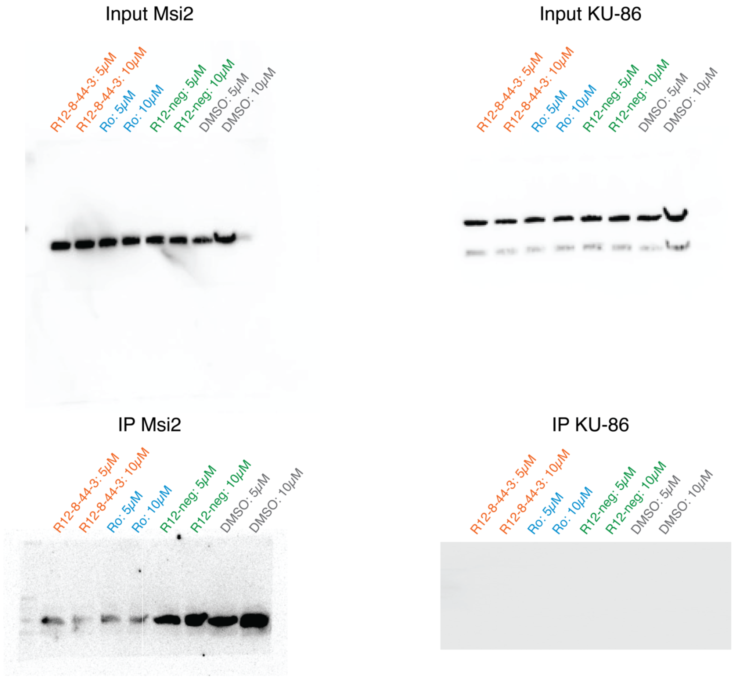


**Figure S8: Uncropped Western blots from pulldown experiment.** Analysis of this experiment is described in **Figure 5**.

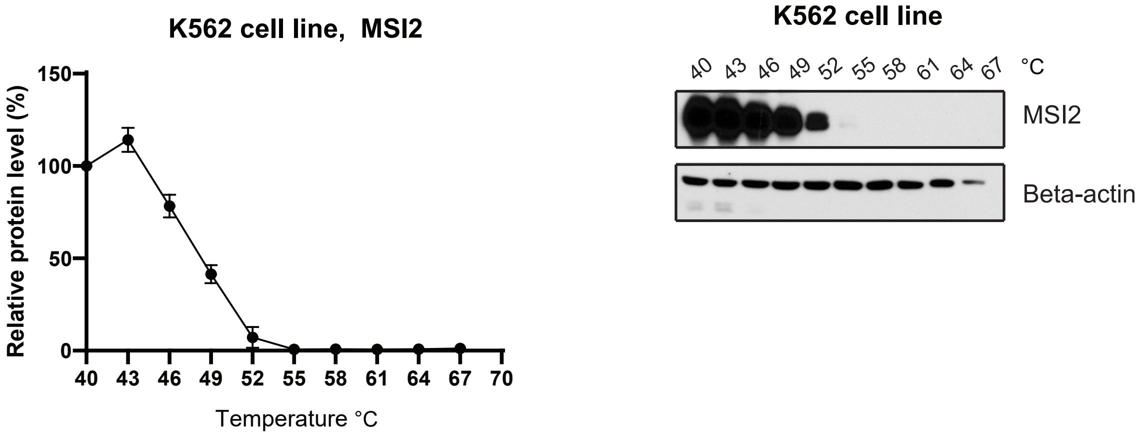

**Figure S9: Thermal-induced aggregation of cellular Msi2 in the K562 cell line.** Cells were briefly incubated at temperatures between 40 °C to 67 °C, then centrifuged and the amount of Msi2 remaining in the soluble fraction was quantified by Western blotting. Data are presented as the average ± S.E.M. from 3 independent replicates, normalized first to beta-actin and then to the data at 40 °C. The observed loss of soluble Msi2 upon heating was similar in K562 cells as in PC9 cells (**Figure 6b**), justifying the use of 49 °C for probing the effect of putative binders in both cell lines.

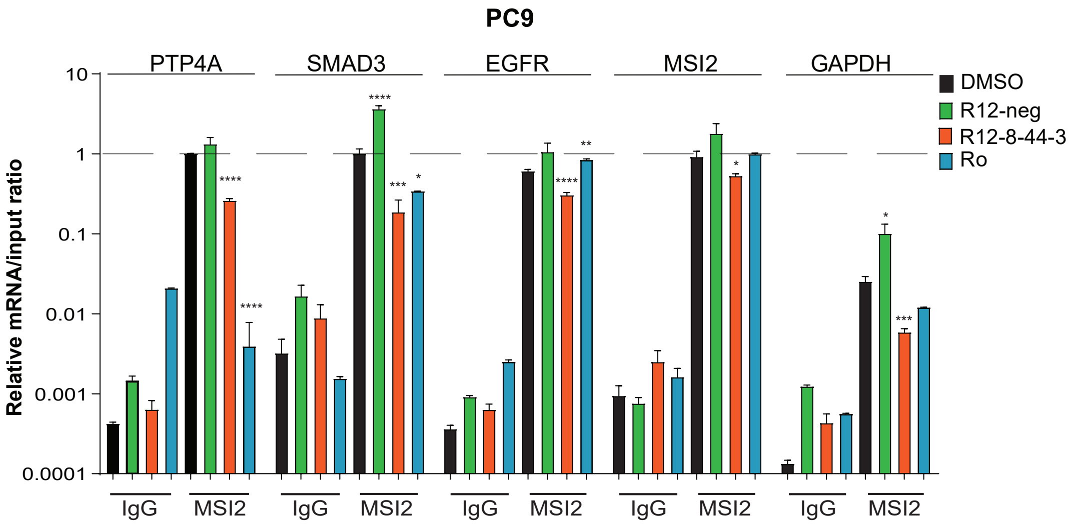


**Figure S10: IgG negative control for mRNA immunoprecipitation (RIP) studies.** PC9 cells were treated with indicated compounds at 10 µM concentrations for 8 hours. PC9 cell lysates were immunoprecipitated using either an anti-Msi2 antibody or a negative control (IgG) antibody, followed by reverse transcription and quantitative RT-qPCR for indicated genes. We emphasize that *data are presented on a log scale*: using IgG instead of anti-Msi2 results in 100x less mRNA for Msi2 targets (as expected). Data are shown as average of inhibition effect of 3 independent experiments. In this plot, * indicates $p<0.05$, ** indicates $p<0.01$, *** indicates $p<0.001$, and **** indicates $p<0.0001$.

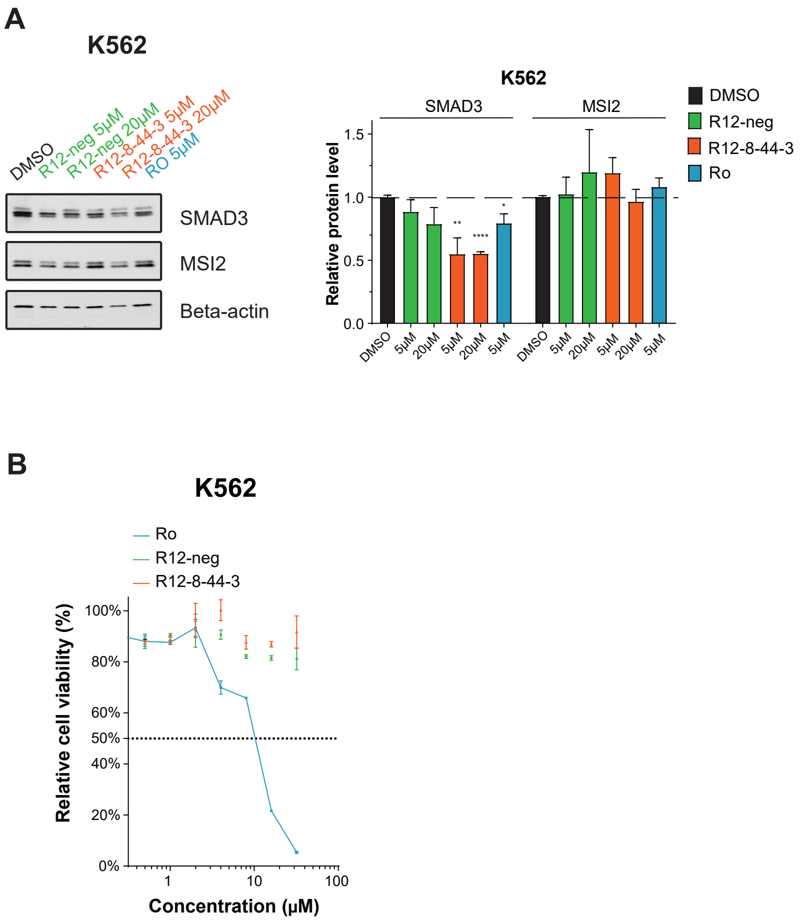


**Figure S11: Effect of R-12-8-44-3 on K562 cell line. (A)** Immunoblot from K562 cells treated with Ro (positive control), R12-neg (negative control), or R12-8-44-3. Cells were treated with indicated compounds at two concentrations (5 µM and 20 µM) for 48 hours, then protein levels of relevant Msi2 targets were determined (EGFR, SMAD3, and Msi2). One representative image is shown; quantification is presented as the average ± S.E.M. from 3 independent replicates, normalized to DMSO-treated cells. Statistical analysis was performed using unpaired two tailed t-test. In this plot, * indicates p<0.05, ** indicates p<0.01, *** indicates p<0.001, and **** indicates p<0.0001. **(B)** Cell viability quantified by CellTiter-Blue of PC9 cell line after 3-day treatment with osimertinib (osi alone), osimertinib with 10 µM R12-8-44-3, or osimertinib with 10 µM R12-neg. Representative data of one of three independent replicates is shown.

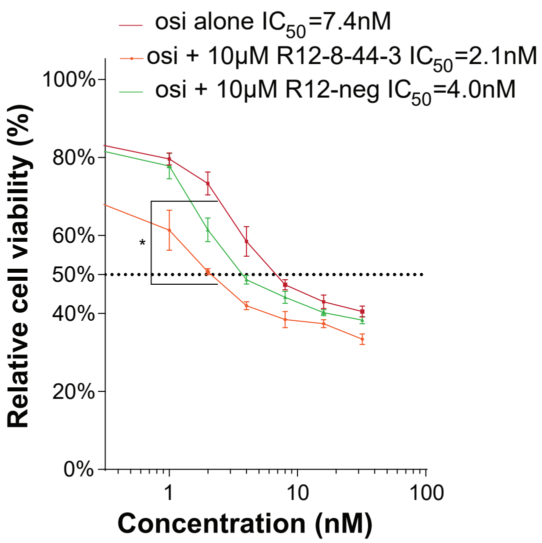


**Figure S12: Sensitization of PC9 cells to osimertinib upon treatment with R-12-8-44-3.** Cell viability was quantified by CellTiter-Blue of PC9 cell line after 3-day treatment with osimertinib (osi alone), osimertinib with 10 µM R12-8-44-3, or osimertinib with 10 µM R12-neg. Representative data of one of three independent experiments is shown. Statistical analysis was performed using one way ANOVA analysis of one of the experiments, $p<0.02$. Error bars indicate S.E.M.
